# RNA-Seq analysis identifies novel roles for the primary cilia gene *SPAG17* and the *SOX9* locus non-coding RNAs in systemic sclerosis

**DOI:** 10.1101/2021.11.20.468677

**Authors:** Elisha D.O. Roberson, Mary Carns, Li Cao, Kathleen Aren, Isaac A. Goldberg, David J. Morales-Heil, Benjamin D. Korman, John P. Atkinson, John Varga

## Abstract

Systemic sclerosis (**SSc**) is characterized by immune activation, vasculopathy, and unresolving fibrosis in the skin, lungs, and other organs. We performed RNA-Seq analysis on skin biopsies and peripheral blood mononuclear cells (**PBMCs**) from SSc patients and controls to better understand SSc pathogenesis. We analyzed these data to 1) test for case-control differences, and 2) identify genes whose expression levels correlate with SSc severity as measured by local skin score, modified Rodnan skin score (**MRSS**), forced vital capacity (**FVC**), or diffusion capacity for carbon monoxide (**DLCO**). We found that PBMCs from SSc patients showed a strong type 1 interferon signature. This signal replicated in the skin, with additional signals for increased extracellular matrix (**ECM**) genes, classical complement pathway activation, and the presence of B cells. Notably, we observed a marked decrease in the expression of *SPAG17*, a cilia component, in SSc skin. We identified genes that correlated with MRSS, DLCO, and FVC in SSc PBMCs and skin using weighted gene co-expression analysis (**WGCNA**). These genes were largely distinct from the case/control differentially expressed genes. In PBMCs, type 1 interferon signatures negatively correlated with DLCO. In SSc skin, ECM gene expression positively correlated with MRSS. Network analysis of SSc skin genes correlated with clinical features identified the non-coding RNAs *SOX9-AS1* and *ROCR*, both near the *SOX9* locus, as highly connected, “hub-like” genes in the network. These results identify non-coding RNAs and *SPAG17* as novel factors potentially implicated in SSc pathogenesis.

## INTRODUCTION

Systemic sclerosis (**SSc**), is a complex orphan disease characterized by autoantibodies, vasculopathy of small vessels, and synchronous / unresolving fibrosis in multiple organs (Allanore et al., 2015, van den Hoogen et al., 2013). There is substantial patient-to-patient heterogeneity in clinical features, disease severity, and the rates of progression. Currently, there are few effective treatments for SSc. Moreover, there is a lack of molecular biomarkers that reliably predict clinical course, reflect disease activity, or identify rational therapeutic targets (Varga and Roberson, 2015).

One approach to improve our understanding of the evolution and progression of the disease is through transcriptomics. Previous primary and secondary analyses of transcriptome data in SSc used microarray, bulk RNA-Seq, and single-cell RNA-Seq approaches. These studies revealed molecular heterogeneity among individual transcriptomes, increased type 1 interferon signaling, potential molecular subtypes, and altered cell populations in the skin (Apostolidis et al., 2018, Assassi et al., 2015, Derrett-Smith et al., 2017, Karimizadeh et al., 2019, Pendergrass et al., 2012, Skaug et al., 2020). We sought to further clarify molecular disruptions in SSc, to find correlations with clinical measures of disease activity, and to determine if expression-trait correlation gene sets overlap with the case/control differential expression gene sets. We used prospective collection of skin and PBMC samples from control subjects and patients with SSc followed by bulk RNA-Seq. At each visit, disease severity was assessed by the local skin score, modified Rodnan skin score (**MRSS**), and pulmonary function testing. For RNA-Seq, we used a ribosomal depletion method to permit detection of both nascent and mature mRNA along with non-coding RNAs lacking a poly(A) tail. This method may be more sensitive for genes with low expression levels or short half-life than poly(A)-based RNA-Seq methods, enabling us to identify potentially overlooked contributors to SSc. These methodologies allowed us to examine categorical differences between SSc patients and unaffected controls, as well as to identify genes whose expression is correlated with alterations in established clinical parameters of disease progression.

## RESULTS

### Study cohort and demographics

Systemic sclerosis cases (n=21) were recruited from the Northwestern Scleroderma Clinic and fulfilled classification criteria for SSc (van den Hoogen et al., 2013). These patients were further classified into limited cutaneous SSc (**lcSSc**; n=5), diffuse cutaneous SSc (**dcSSc**; n=14), SSc sine scleroderma (**SSS**; n=1), and very early diagnosis of systemic sclerosis (**VEDOSS**; n=1). Controls were volunteers with no history of an autoimmune or inflammatory disease (n=14). At each study visit, we obtained a whole-blood sample and two 3 mm skin punch biopsies. We also obtained pulmonary function tests, and the same observer assessed the MRSS and local skin score. Seven dcSSc, two lcSSc, and one SSS individual volunteered for a second sampling and assessment at a 6-month follow-up visit (**Table 1**).

**Table 1.**
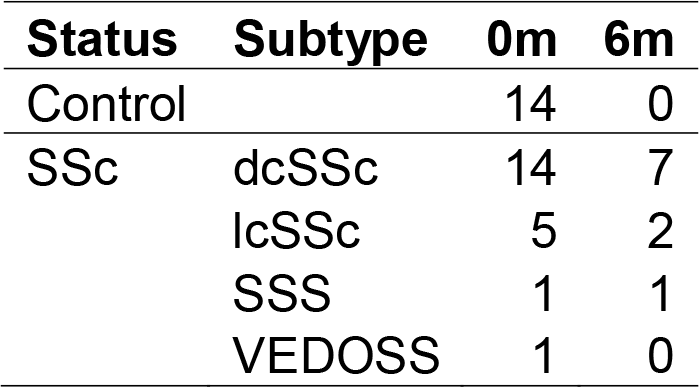
Sample subtype counts. Each time point collected consisted of both a blood sample and a skin punch biopsy. Controls were only sampled at enrollment. Some systemic sclerosis patients were re-sampled approximately 6 months after enrollment. A total of 21 systemic sclerosis patients were enrolled in the study, and 10 opted to have the secondary sample. For general differential expression between classes, we used only the 0m samples. For correlation with disease activity, we used all samples. Abbreviations: SSc, systemic sclerosis; dcSSc, diffuse cutaneous SSc; lcSSc, limited cutaneous SSc; SSS, systemic sclerosis sine scleroderma; VEDOSS, very early diagnosis of systemic sclerosis.

| Status | Subtype | 0m | 6m |
| --- | --- | --- | --- |
| Control |  | 14 | 0 |
| SSc | dcSSc | 14 | 7 |
|  | lcSSc | 5 | 2 |
|  | SSS | 1 | 1 |
|  | VEDOSS | 1 | 0 |

For group-wise demographic summaries, we included controls and subjects with either lcSSc or dcSSc (**Table 2**). Control subjects were significantly younger than SSc patients. Within the SSc cohort, the lcSSc and dcSSc subsets were balanced for age and disease duration (**Table 3**). There were more women in the dcSSc than the lcSSc subset, while self-declared ethnicity was similar between the two.

**Table 2.** Case/control demographics. Basic demographic information for our study cohort. All values are “n (%)” unless otherwise indicated. The controls were individuals without a self-reported history of autoimmune disease. SSc met ACR criteria for a definitive diagnosis. The controls were significantly younger than the SSc cases (two-sided student t-test) but balanced for sex (Fisher’s exact test). The control and SSc cohorts were predominantly white, and the population background was significantly different between the two (Fisher’s exact test). Abbreviations: SSc, systemic sclerosis; SD, standard deviation.

|  | **Control (n=14)** | **SSc (n=19)** | **P-value** |
| --- | --- | --- | --- |
| **Age mean (SD)** | 32.6 (10.8) | 49.7 (11.4) | $1.30 \times 10^{-4}$ |
| **Female** | 9 (64.3) | 15 (78.9) | $4.42 \times 10^{-1}$ |
| **Ethnicity** | | | $7.22 \times 10^{-3}$ |
| **Asian** | 0 (0.0) | 2 (10.5) |  |
| **Black** | 3 (21.4) | 2 (10.5) |  |
| **Hispanic** | 5 (35.7) | 0 (0.0) |  |
| **White** | 6 (42.9) | 15 (78.9) |  |

**Table 3.** Systemic sclerosis subtype demographics. Shown are the basic demographic and antibody staining pattern information for the two main systemic sclerosis sub-groups: diffuse disease and limited disease. Values in the table are represented as “mean (standard deviation)” unless otherwise indicated as “n (%)”. We determined the significance of count data (sex, ethnicity, immunofluorescence) using Fisher’s exact test. We calculated the significance of continuous data (age, duration, MRSS, body mass index, lung parameters) using a two-sample t-test with unequal variance. The MRSS, body mass index, forced vital capacity, diffusion capacity for CO, and total lung capacity were calculated at repeat visits as well. For each individual, we used the “worst” observed value as the representative value. Abbreviations: MRSS, Modified Rodnan Skin Score; BMI, body mass index; FVC, forced vital capacity; DLCO, diffusion capacity for carbon monoxide; TLC, total lung capacity. *These lung function parameters were calculated as percent estimated maximum for age and sex. †Immunofluorescence data was only available for 12/14 dcSSc individuals. Percents were calculated based on this availability.

|  | **dcSSc (n=14)** | **lcSSc (n=5)** | **P-value** |
| --- | --- | --- | --- |
| **Age mean** | 49.7 (11.7) | 49.8 (11.9) | $9.89 \times 10^{-1}$ |
| **Female (%)** | 13 (92.9) | 2 (40.0) | $3.74 \times 10^{-2}$ |
| **Mean duration, months** | 42.7 (33.2) | 36.2 (22.8) | $6.39 \times 10^{-1}$ |
| **Deaths (%)** | NA | NA |  |
| **MRSS** | 24.1 (10.2) | 8.8 (5.0) | $5.93 \times 10^{-4}$ |
| **Forearm MRSS** | 1.9 (0.9) | 0.8 (0.8) | $3.66 \times 10^{-2}$ |
| **BMI** | 26.7 (5.5) | 29.5 (6.3) | $4.02 \times 10^{-1}$ |
| **FVC\*** | 75.1 (15.6) | 82.6 (12.5) | $3.14 \times 10^{-1}$ |
| **DLCO corrected\*** | 66.2 (20.8) | 80.0 (17.4) | $1.88 \times 10^{-1}$ |
| **TLC\*** | 85.6 (17.2) | 85.8 (11.7) | $9.83 \times 10^{-1}$ |
| **Ethnicity (%)** | | | $7.42 \times 10^{-1}$ |
| **Asian** | 1 (7.1) | 1 (20.0) |  |
| **Black** | 2 (14.3) | 0 (0.0) |  |
| **Hispanic** | 0 (0.0) | 0 (0.0) |  |
| **White** | 11 (78.6) | 4 (80.0) |  |
| **Immunofluorescence† (%)** | | | $4.29 \times 10^{-1}$ |
| **Centromere** | 1 (8.3) | 2 (40.0) |  |
| **Homogeneous** | 3 (25.0) | 0 (0.0) |  |
| **Nucleolar** | 4 (33.3) | 1 (20.0) |  |
| **Speckled** | 7 (58.3) | 3 (60.0) |  |

### Including intronic read counts in the ribosomal depletion library data improves quantification

We were interested in detecting transcripts that may be in a minority of cells or have a short half-life. For this reason, we used stranded, ribosomal depletion library kits to detect polyadenylated mature mRNA, unspliced mRNA, and long non-coding RNA. We used the Picard tools RNA-Seq metrics to determine if the stranded prep worked, and what fraction of mapped bases overlapped different types of genomic regions (**Fig. 1a**). The strandedness did perform well, with an average of 96.4% (± 1.1% standard deviation [**SD**]) of mapped bases overlapping a known gene on the correct strand. With regard to what genomic features reads mapped to, a small portion of mapped bases was in intergenic regions (8.8% ± 3.1%) or ribosomal RNA (5.0% ± 1.8%). The low fraction mapping to ribosomal sources indicates that the ribosomal depletion worked well. A greater fraction of mapped bases (19.0% ± 1.8%) were in coding exons. However, most of the mapped bases were within UTRs (16.6% ± 1.1%) and introns (50.6% ± 3.2%). By focusing on coding exons or even UTRs + coding exons, we would only be considering approximately 35.6% of mapped bases. Given the (sometimes substantially) larger size of introns versus exons, this is not necessarily surprising. We compared the normalized per-sample abundance estimates (regularized logarithm) using the GTF-guided counts (exon overlapping reads only) to the total gene read counts (UTR + coding exons + introns; **Fig. 1b-c**). For the total gene reads, we counted any read that overlapped with a known gene on the correct strand. The linear regression of the normalized data for exon-overlapping versus all reads correlated reasonably well overall (adjusted R^2^ = 0.8605), but the exon-only method underestimated abundance due to having fewer usable counts (**Fig. 1b** estimated exon-only coefficient 0.911 of the all read abundance, P < 2.2 x 10^-16^). This effect would be amplified for low-expression genes that might have few coding-exon reads compared to intronic reads. We, therefore, performed all differential expression and correlation analyses using the total gene read counts.

**Fig. 1.**
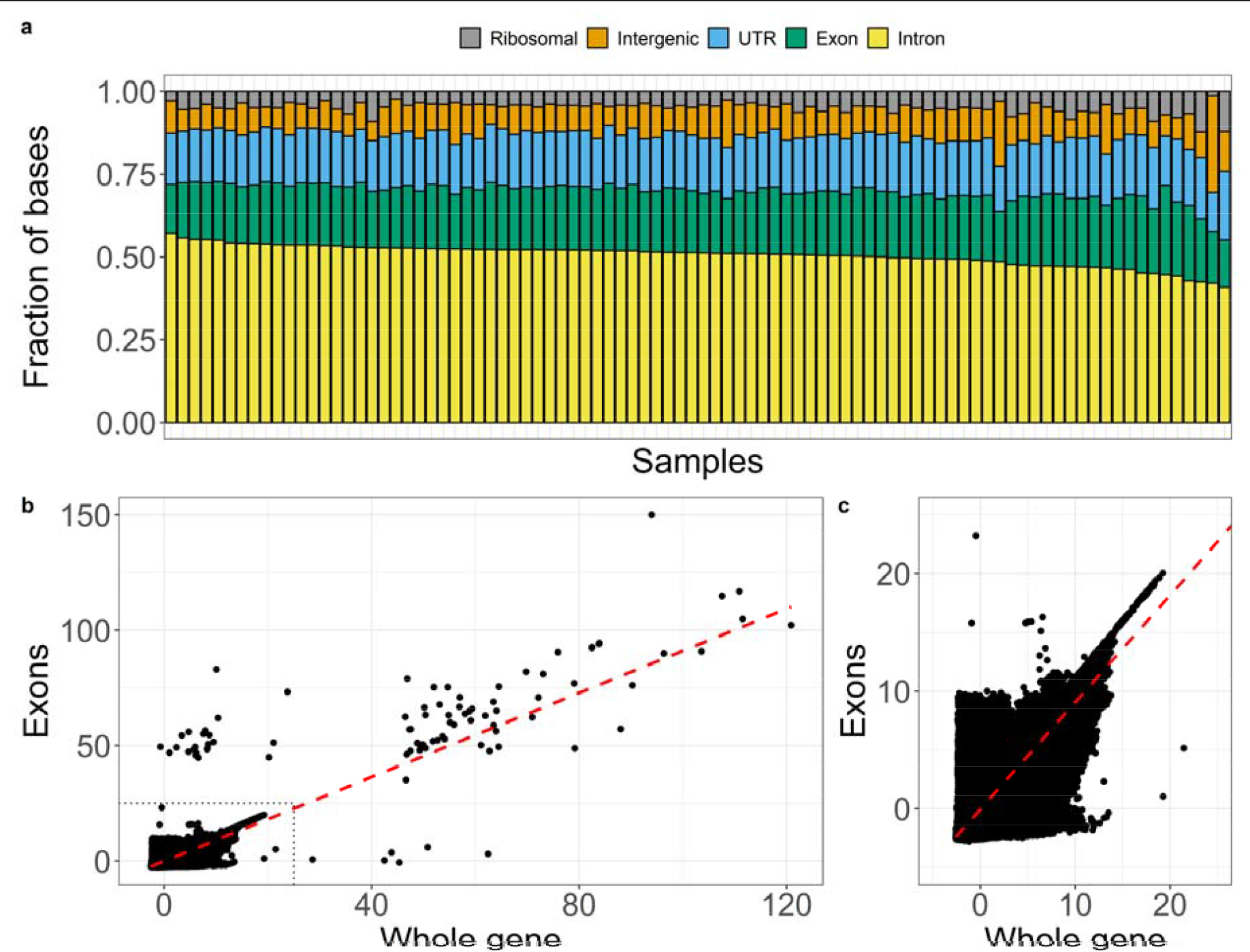
Ribosomal depletion library sequenced bases primarily map to introns. **a.** Individual samples are shown on the x-axis, and the fraction of mapped bases of different classes (ribosomal RNA, intergenic bases, gene untranslated regions [**UTR**], coding exonic bases, and intronic bases) on the y-axis. The fraction of mapped bases in each category is shown as a stacked bar chart. Most of the ribosomal RNA was successfully removed. Most of the sequenced bases were derived from intronic regions. **b.** The x-axis is the regularized logarithm (normalized expression) using all counts overlapping a gene, and the y-axis shows the normalized expression using counts derived from exon overlapping reads only. The red dashed line is the best fit using linear regression. **c.** Shows a zoomed view of the grey dashed line in the bottom-left corner of **b**. Both **b** & **c** show that the estimated abundance somewhat correlates between the two strategies. The linear regression coefficient estimated that the exon-only method generally underestimated abundance compared to all gene overlapping reads.

### SSc PBMCs demonstrate evidence of increased type 1 interferon signaling

We sought to characterize gene expression changes of SSc PBMCs since it is a minimally invasive tissue source. For cases, we only included baseline lcSSc and dcSSc samples to avoid bias toward individuals sampled more than once. There were 147 genes with decreased expression in SSc PBMCs (113 at least -1.5-fold) and 100 genes with increased expression (61 at least 1.5-fold; **Fig. 2a; Table ST1**). The most significantly decreased genes included *GALNTL6* (polypeptide N- aceltylgalactosaminyltransferase-like 6; -4.09 fold-change [**FC**]), *GPM6A* (glycoprotein M6A; -3.98 FC), *SLC4A10* (solute carrier family 4 member 10; -3.33 FC), *COL4A3* (collagen type IV alpha 3 chain; -4.36 FC), *OSBPL10* (oxysterol-binding protein-like 10; -2.88 FC), and *NRCAM* (neuronal cell adhesion molecule; -3.35 FC). The PBMCs of individuals with sporadic Meniere’s disease have decreased *SLC4A10* (Sun et al., 2018). *BANK1* (B Cell Scaffold Protein with Ankyrin Repeats 1) was decreased -2.58-fold in SSc. It is also decreased in the peripheral blood of mice using the collagen-induced arthritis model (Yang et al., 2018). We next tested for enrichment of known molecular pathways among genes with decreased expression in SSc (**Fig. 2b**; **Table ST2**). The most consistent trend was enrichment for collagen pathways due to decreases in *COL4A3* and *COL4A4*.

**Fig. 2.**
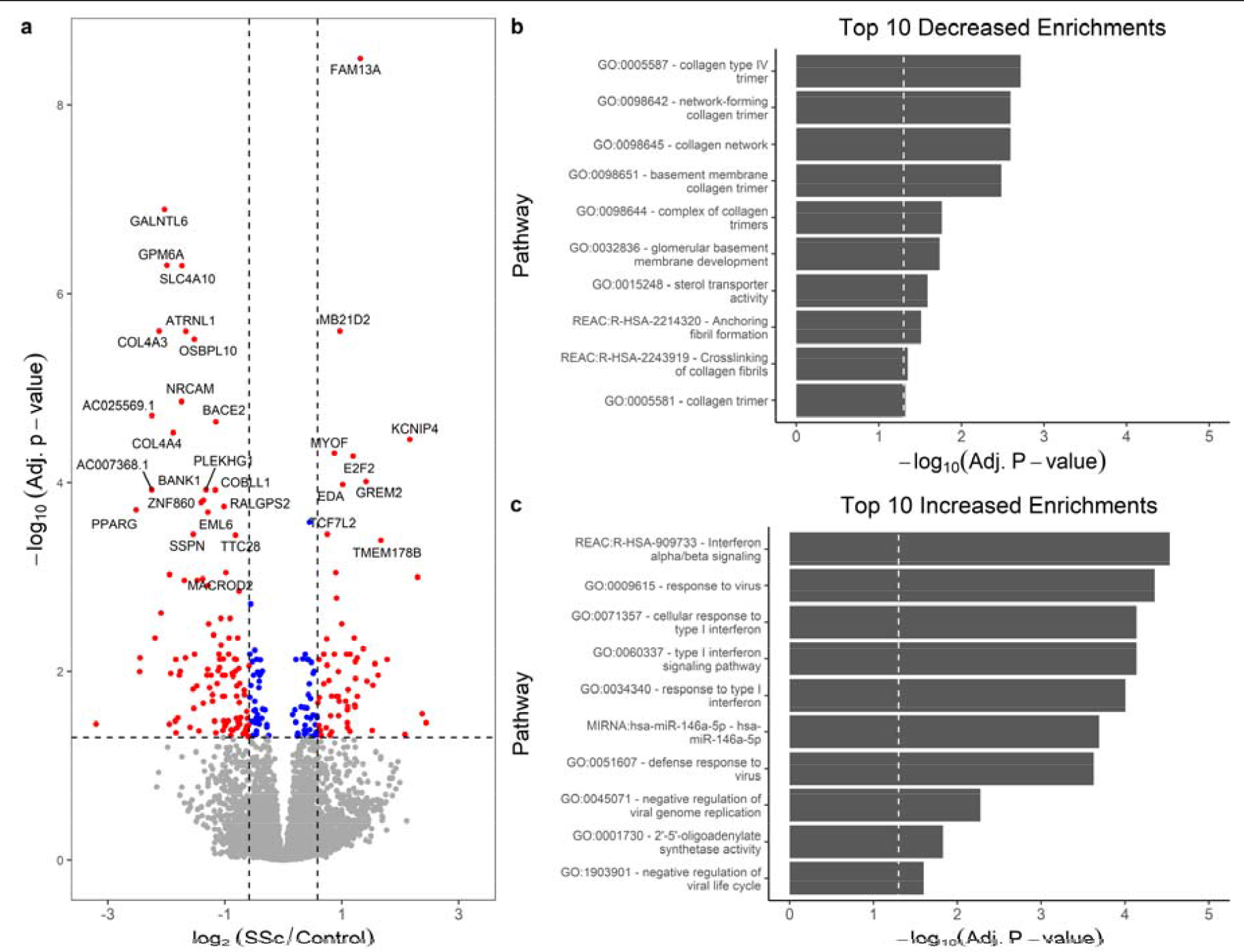
Systemic sclerosis PBMCs have strong enrichments for increased type 1 interferon signaling. **a.** Standard volcano plot. The x-axis shows the effect size on a log2 scale of systemic sclerosis over control. The y-axis is the significance of the difference (-log10 scale). The vertical dashed lines show the 1.5-fold cutoff. The horizontal line is the 0.05 adjusted significance threshold. Genes meeting a 1.5-fold cutoff and transcriptome-wide significance are in red. Significant points that don’t meet the fold-change cutoff are blue. Genes that weren’t significantly different are in grey. **b.** Shows the enriched pathways for genes decreased in SSc PBMCs. The x-axis shows the -log10 of the adjusted p-values. The dashed vertical line shows the 0.05 adjusted p-value cutoff. The y-axis shows the pathway. The most significant enrichments include collagen and sterol transporter. **c.** The axes are the same as in **b**, but the pathways tested were for genes at least 1.5-fold increased in SSc PBMCs. The predominant signal is increased type 1 interferon signaling.

Genes with significantly increased expression in SSc PBMCs compared to controls included *FAM13A* (family with sequence similarity 13, member A; 2.14), *E2F2* (E2F transcription factor; 2.18), *MYOF* (myoferlin; 1.76), and *FNIP2*(folliculin interacting protein 2; 1.37). SNPs in *FAM13A* are associated with an increased risk of pulmonary fibrosis (Fingerlin et al., 2013) and liver cirrhosis (Zhang Y. et al., 2019). *E2f2* is required for myeloid cell development in mice (Trikha et al., 2011) and is a regulator of inflammatory signaling via interactions with NF-κB (Ankers et al., 2016, Wang et al., 2018). Myoferlin chaperones phosphorylated STAT3 to the nucleus, mediating IL6/STAT3 signaling (Yadav et al., 2017).

We tested the genes with increased expression in SSc PBMCs for pathway enrichment (**Fig. 2c; Table ST3**). The most significant pathways were related to type 1 interferon activity, including interferon-alpha/beta signaling (Reactome R-HSA-909733), response to type 1 interferon (GO:0071357), and type 1 interferon signaling (GO:0060337). These enrichments were driven by increases in the expression of classic interferon-responsive genes, including *IFIT1*, *IFIT3*, *IFITM3*, *OAS1*, *OAS3*, and *RSAD2*. Targets of two transcription factors were significantly enriched: IRF5 (TransFac M04016) and IRF9 (M11680). *IRF5* is highly expressed in M1 polarized human macrophages (Krausgruber et al., 2011). A high prevalence of M1-like macrophage signatures has been detected in SSc skin (Skaug et al., 2020).

We observed a robust increase in type 1 interferon signatures in SSc PBMCs. Previous studies have observed similar increased expression of type 1 interferon-stimulated genes in SSc PBMCs as well as in skin (Assassi et al., 2010, Duan et al., 2008). There is evidence that the increased type 1 interferon response general and the increased IRF7, in particular, may play a direct role in increasing fibrosis in SSc (Wu et al., 2019). It is also interesting to consider whether some of these signatures, including the increase in IRF5 and its known high expression in M1-polarized macrophages, indicate a specific role for this macrophage subpopulation in systemic sclerosis.

### SSc skin shows evidence of substantial immune activation, increased complement component **expression, and loss of ciliary protein *SPAG17***

PBMC procurement is minimally invasive, but may not reflect the molecular biology of affected tissues. The skin may therefore provide more insight into the molecular defects in SSc. We again compared controls to all the lcSSc and dcSSc baseline biopsies. There were 526 genes significantly decreased (226 at least -1.5-fold) and 1,200 genes increased (816 at least 1.5-fold; **Fig. 3a**; **Table ST4**) in SSc compared to control skin biopsies. There are nearly seven times as many differentially expressed genes in the skin biopsy comparison versus the PBMC comparison, highlighting the advantages of studying affected tissue.

**Fig. 3.**
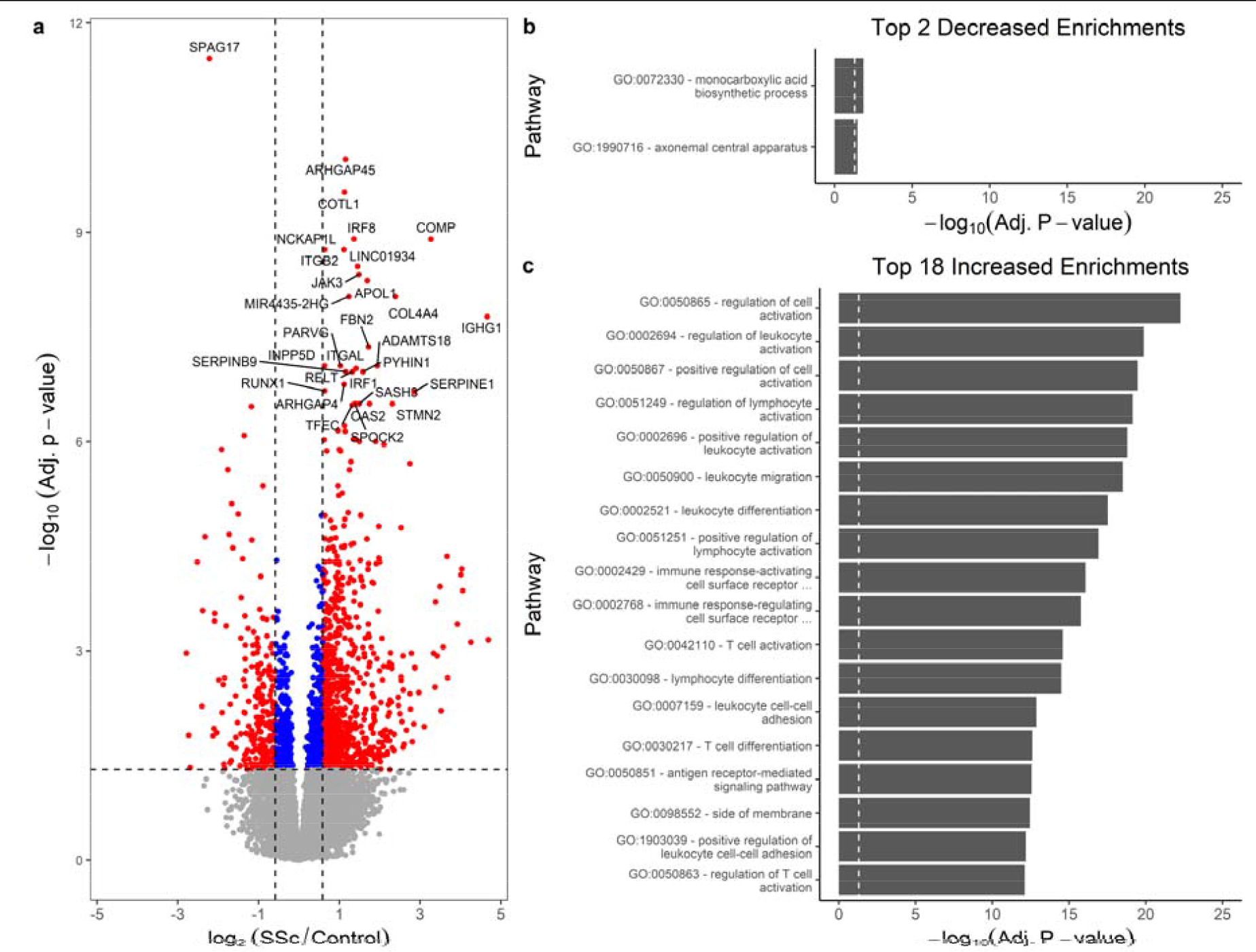
SSc skin has decreased primary cilia protein *SPAG17* and increased immune activation. **a.** Volcano plot showing the significance and effect-sizes of gene expression in systemic sclerosis versus control skin. The log2 fold-change is on the x-axis, and the -log10 adjusted significance is on the y-axis. The vertical lines indicated 1.5-fold up and down cutoffs. The horizontal line shows the 0.05 adjusted significance threshold. The most significant difference was a decrease in *SPAG17* in SSc skin. More genes were significantly increased in SSc than significantly decreased. Both **b.** and **c.** show the gProfilerR pathway enrichments. The x-axis shows the -log10 of the significance for each enrichment. The y-axis shows the name of the enriched pathway. **b.** There were few significant enrichments for decreased genes, including the ciliary axoneme. **c.**The most significant enrichments for genes increased in SSc were related to immune cell activation, indicating the migration of immune cells into the skin.

The most significantly altered gene in the entire study was a decrease of *SPAG17* in SSc skin samples (sperm-associated antigen 17; -4.67 FC; adjusted p-value 3.22 x 10^-12^). To our knowledge, this gene had not been previously associated with differential expression in systemic sclerosis skin. Its expression level is relatively low, and studies using hybridization microarrays may not have been sensitive enough to detect its expression versus the background fluorescence. The low expression might also cause it to be filtered out of some RNA-Seq studies. SPAG17 protein is required for the function of primary cilia and male fertility (Kazarian et al., 2018). Mice deficient in *Spag17* have bone abnormalities such as decreased femur length and disrupted femur morphology (Teves et al., 2015). Its role in skin and immune cells is not particularly clear, though as part of the primary cilia it could be involved in signaling. The decreased expression of *SPAG17* does not require appreciable fibrosis. When all SSc subtypes (including baseline and follow-up biopsies) are examined, VEDOSS has control-like *SPAG17* expression. SSc sine scleroderma skin, however, has decreased *SPAG17* expression levels (comparable to lcSSc and dcSSc) despite the lack of skin fibrosis (**Fig. 4**). The similarity of the skin transcriptomes from patients with SSS, lcSSc, and dcSSc is also observed by principal components analysis of the top case/control differentially expressed genes (supplemental material and **Fig. S2**).

**Fig. 4.**
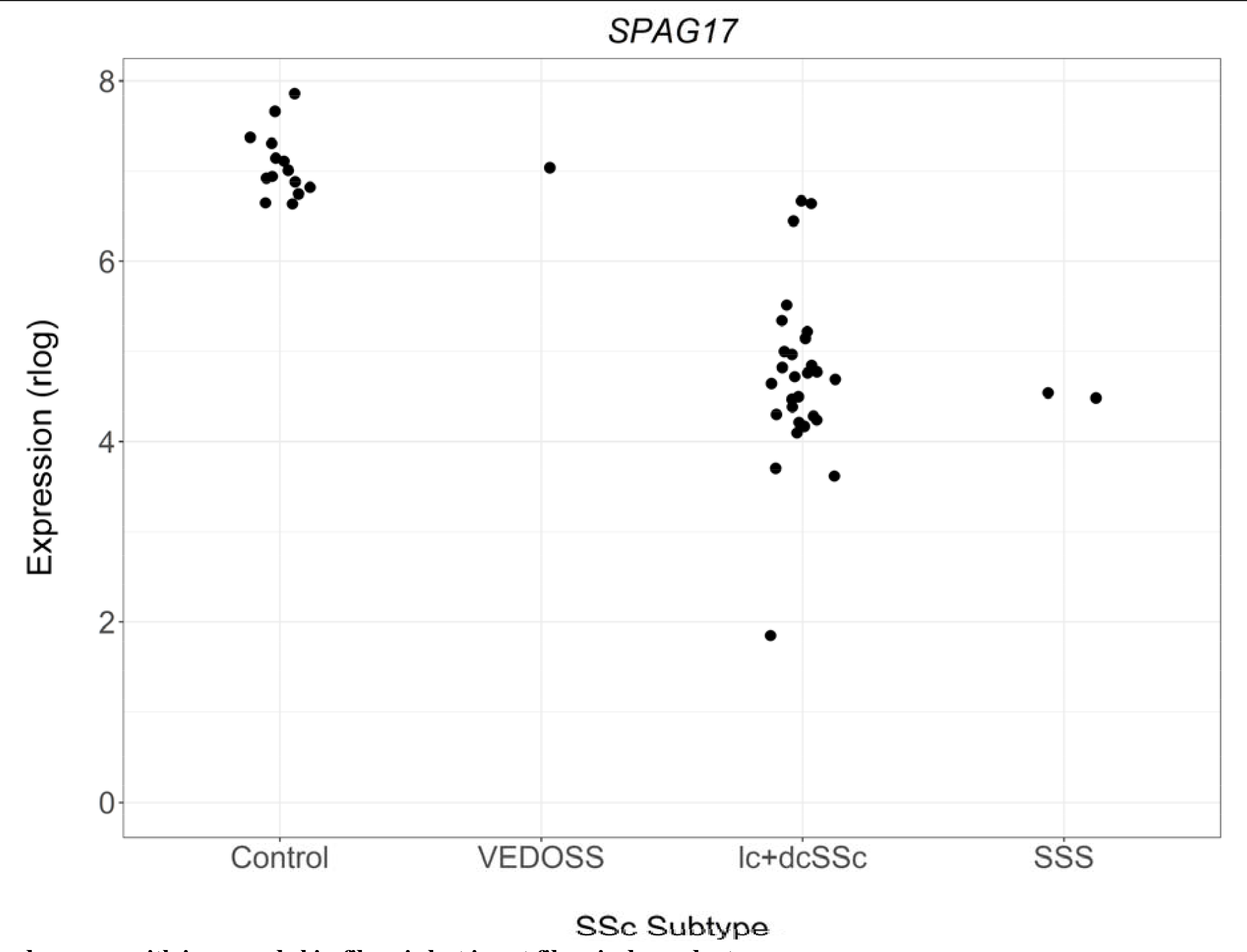
*SPAG17* expression generally decreases with increased skin fibrosis but is not fibrosis dependent Shown are the subtype of SSc (x-axis) and normalized expression (regularized logarithm; y-axis) of *SPAG17* in controls and all SSc samples. Several patterns emerge. Control samples have a relatively limited range of expression of *SPAG17*. The VEDOSS sample, which would not have extensive fibrosis, has a control-like expression. Some of the lcSSc samples have expression that is close to the lowest expressing controls. Most lcSSc and dcSSc samples have decreased expression. The SSS initial and follow-up samples both have SSc-like expression. This suggests that while *SPAG17* is reduced in the majority of SSc samples, it must not be purely a proxy for enhanced fibrosis, because if that was the case SSS samples should have control-like expression. Abbreviations: SSc, systemic sclerosis; SSS, SSc sine scleroderma; VEDOSS, very early diagnosis of systemic sclerosis; lcSSc, limited cutaneous SSc; dcSSc, diffuse cutaneous SSc.

Other genes showing decreased expression included *SEMA3E* (semaphorin 3E; -2.25 FC), *LGR5* (leucine-rich repeat-containing G-protein coupled receptor 5; -2.70 FC), and *TSPAN8* (tetraspanin-8; -2.53 FC). *SEMA3E* functions in the immune system by inhibiting the migration of neutrophils and natural killer cells (Alamri et al., 2018, Movassagh et al., 2017), and by inhibiting the interaction of dendritic cells and thymocytes (Ueda et al., 2016). A decrease in *SEMA3E* might allow for a greater influx and more interactions among immune cells in the skin.

*LGR5* is a member of the G protein-coupled receptor family that is an important target and modulator of Wnt/ β-catenin signaling. Notably, it may be a marker for stem cells in multiple tissues (de Lau et al., 2014). *LGR5* expression is a known marker of intestinal crypt stem cells in mice. It has also recently been shown to be a marker for intestinal villi tip telocytes that maintain the correct differentiation gradient on the villus axis via non-canonical Wnt signaling (Bahar Halpern et al., 2020). Dermal telocytes are lost in fibrotic skin and internal organs in SSc (Manetti et al., 2013, Manetti et al., 2014). If these dermal telocytes also help to coordinate differentiation and/or signaling in the skin, their loss (perhaps detected by the reduction of *LGR5*) may directly contribute to the development of SSc fibrosis.

In the human T1C3 melanoma cell line, knockdown of *Tspan8* causes increased adherence to extracellular matrix (**ECM**) proteins (El Kharbili et al., 2017). The decrease in SSc skin may indicate increased binding to the skin matrix. Only two known pathways were enriched among decreased genes (**Fig. 3b; Table ST5**): monocarboxylic acid biosynthesis (GO:0072330) and axonemal central apparatus (GO:1990716). It’s worth noting that the ciliary axoneme pathway enrichment was only due to the decrease in *SPAG17*.

Genes increased in SSc included *ARHGAP45* (rho GTPase activating protein 45; 2.22 FC), *COTL1* (coactosin-like protein 1; 2.18 FC), *IRF8* (interferon regulatory factor 8; 2.57 FC), and *COMP* (cartilage oligomeric matrix protein; 9.63 FC). The protein translated from *ARHGAP45* is referred to as HA-1 or HMHA1, i.e. the minor histocompatibility protein HA-1. HMHA1 is a negative regulator of endothelial integrity (Amado-Azevedo et al., 2018). The increase in HMHA1 might therefore cause increased vascular permeability. COTL1 protein localizes to the immune synapse in T cells after stimulation with CD28 and T cell receptor, which may indicate the presence of stimulated T cells in the skin (Kim et al., 2014). In mouse bone marrow-derived macrophages, Irf8 drives expression of *Naip2*, *Naip5*, *Naip6*, and *Nlrc4*, and is required for optimal activation of the Nlrc4 inflammasome (Karki et al., 2018). The human ortholog of *Nlrc4* (sometimes called *CARD12*) was also increased in SSc skin (*NRLC4*; NLR family CARD domain containing 4; 2.34 FC). The thrombospondin COMP is known to be increased in multiple fibrotic skin conditions (Agarwal et al., 2013).

The extensive list of genes increased in SSc skin would allow us to posit interesting hypotheses for almost any one of them. We, therefore, chose to check these genes for the enrichment of known molecular pathways as well. We found a total of 366 significant enrichments (**Fig. 3c**; **Table ST6**): 1 from CORUM (Giurgiu et al., 2019), 241 Gene Ontology (**GO**) biological pathways (The Gene Ontology Consortium, 2019), 31 GO cellular components, 11 GO molecular functions, 8 from KEGG (Kanehisa and Goto, 2000), 21 from Reactome (Jassal et al., 2020), 44 TRANSFAC transcription factors (Wingender et al., 1996), and 9 WikiPathways (Kutmon et al., 2016). There was enrichment of targets of the transcription factors IRF2, IRF4, IRF5, IRF7, IRF8, IRF9, and ISGF3. The RNA expression of transcription factors *IRF1* (2.65 FC), *IRF5*(1.48 FC), *IRF7* (1.99 FC), and *IRF8*(2.57 FC) were all significantly increased in the skin as well.

Some of the enrichments were variations on similar themes. One of these was the infiltration of immune cells into the skin, including categories for adhesion (GO:0007159, GO:1903039, GO:0033627) and migration (GO:0050900, GO:0002687, GO:0036336, GO:1902624). The genes most frequently associated with the adhesion categories included *ITGB2*, *JAK3*, *RUNX1*, *CD274* (*PD-L1*), and *CD86*. For migration, it included *CCL19*, *CCL5* (*RANTES*), *CCR7*, *CXCL10*, *RAC2*, and *CD74*. Other immune enrichments were related to immune cell differentiation, including dendritic cells (GO:0002573, GO:0097028). Increased plasmacytoid DC infiltrate has been previously observed in SSc skin and those cells secrete interferon-alpha and CXCL4 (Ah Kioon et al., 2018). There was an enrichment of B cell activation and proliferation genes (GO:0042113, GO:0050871, GO:0030890), suggesting that class-switched B cells may be present in the skin. It has been previously shown that direct cell-cell contact between B cells and systemic sclerosis fibroblasts increases excretion of IL-6 and collagen, along with a concurrent increase of alpha smooth muscle actin in the fibroblasts (Francois et al., 2013).

The most obvious feature of affected SSc skin is the increased deposition of connective tissue.

Consistent with this observation, pathways related to ECM and matrix organization were enriched (GO:0031012, GO:0062023, KEGG:04514, REAC:R-HAS-1474244) because of increased expression of *ICAM1*, *AGRN*, *COL10A1*, *COL4A3*, *COL4A3*, *COL4A4*, *COMP*, *CTSS*, and *EMILIN1*.

Interestingly, there was enrichment for pathways related to complement activation, mostly via the classical (antibody) pathway (GO:0006958, GO:0006956, GO:0030449). This enrichment was due to differential expression of immunoglobulin and complement genes (including *C1QB*, *CR1*, *C5AR1*, and *C7*). Complement activation usually leads to the assembly of the membrane attack complex (**MAC**) that can insert into membranes and lyse cells. The MAC is composed of C5b, C6, C7, C8, and C9. MAC fragments are deposited in the dermal vasculature of systemic sclerosis patients, supporting a role for antibody-induced complement activation in SSc vasculopathy (Scambi et al., 2015). These data suggest that aberrant activation of complement may partially mediate cutaneous tissue damage in SSc.

### Transcriptome alterations in SSc skin do not overlap with changes in PBMCs

Differentially expressed genes between conditions can be due to alterations of the transcriptional profile of resident cells, changes in the local cell population (such as an influx of immune cells), or a combination of these factors. If the primary change in SSc skin is the influx of immune cells, we might find extensive overlap in the differentially expressed genes of skin and PBMCs. That does not appear to be the case for SSc. Most differentially expressed genes were unique in the skin (1,018 genes) or PBMC (150 genes) sets (**Fig. 5**). Only 24 genes were differentially expressed in both tissues (**Table ST7**).

**Fig. 5.**
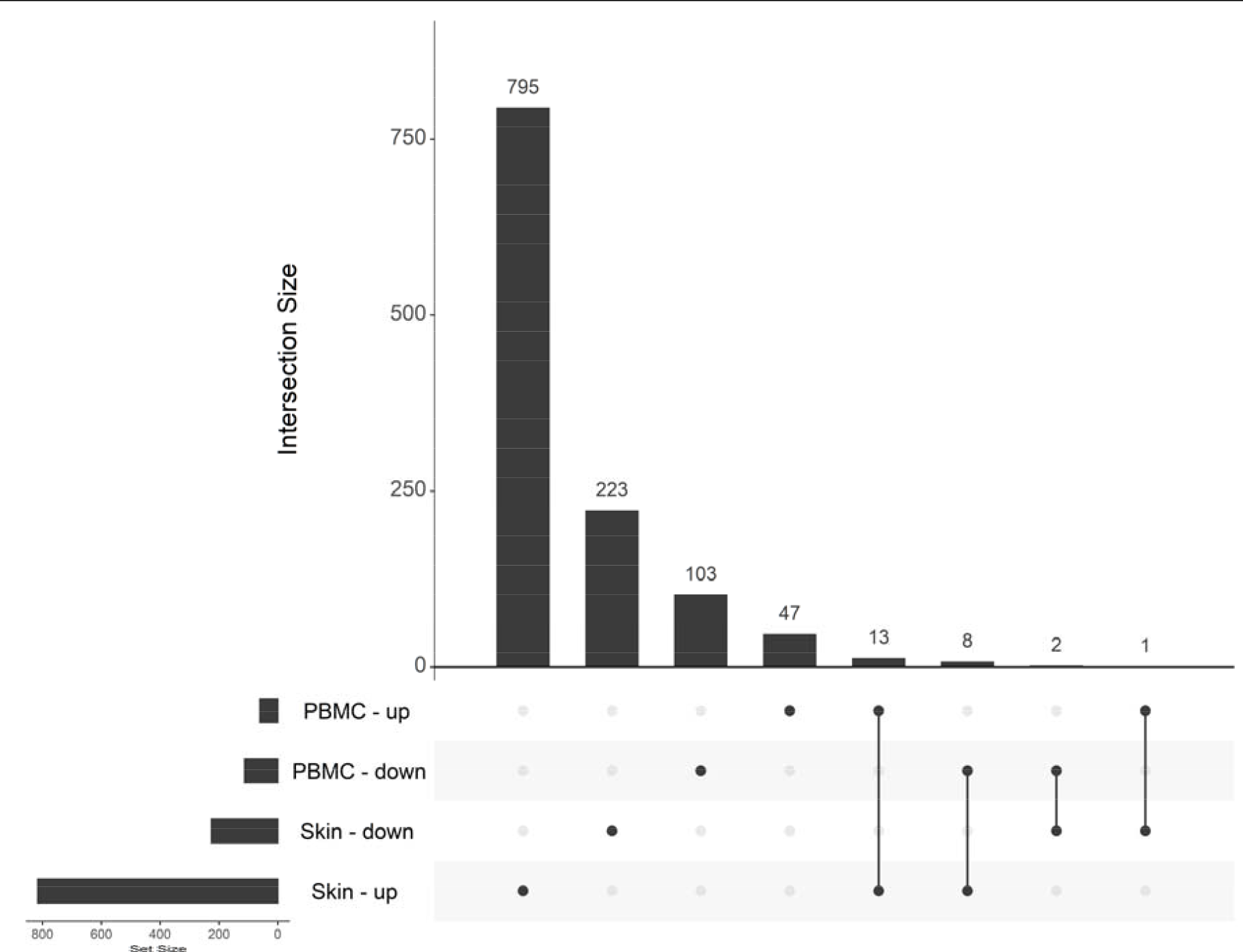
SSc skin and PBMCs have different sets of differentially expressed genes. UpSet chart showing the intersections between differentially expressed genes in the skin and PBMCs. There are 4 possible categories: increased in SSc skin, decreased in SSc skin, increased in SSc PBMCs, and decreased in SSc PBMCs. The Set Size (lower left) shows the total number of genes in each of those categories. Each intersection is represented as dots connected by solid lines (lower section). The Intersection size (upper section) shows the number of genes within that particular intersection. Most DE genes were unique for each tissue, though there were some concordant and discordant overlaps.

There were 15 genes with concordant differential expression, either increased (13) or decreased (2) in SSc for both tissues. As one might hypothesize, several interferon genes (*IFI44*, *IFI44L*, *IFITM3*, *OAS1*, *OAS3*, and *RSAD2*) were concordantly changed in SSc skin and PBMCs.

Interestingly, there were nine genes with significant fold-changes in both tissues but in opposing directions. *GALNT13* was increased in SSc PBMCs (2.96 FC) but decreased in SSc skin (-2.84 FC). The remaining eight genes had decreased expression in PBMCs, but increased expression in skin: *ADAM12*, *COL4A3*, *COL4A4*, *IFNG-AS1*, *IRAK2*, *RUBCNL*, *SEMA6B*, and *SLC4A10*. We cannot tell, based on these data, if this discordance is due to discordant regulation in different cell types of the skin and PBMCs, are markers of the translocation of a specific immune subset from the blood to the skin, or are a response to moving into a proinflammatory/profibrotic environment.

### Gene expression in SSc PBMCs and skin correlate with skin fibrosis and lung functions but are **mostly missed by differential expression analysis**

Group-wise gene expression analysis categorizes samples as test (SSc) or reference (control). This is a reasonable way to determine general features of a trait but ignores the heterogeneity of phenotypes within the trait. SSc in particular has substantial clinical heterogeneity. We used weighted gene co-expression network analysis (**WGCNA**) to correlate gene expression with skin severity (MRSS and forearm local skin score) and lung function (FVC and DLCO). We included all SSc samples for which there was a matching MRSS, FVC, or DLCO measurement for the visit, i.e. follow-ups were included as separate measurements. The tally of positive and negative correlations is listed in **Table 4**.

**Table 4.** Correlation with disease severity parameters. The table shows the type of gene-trait correlation (negative or positive) and counts for each tissue and clinical trait. PBMC genes most often correlated with MRSS, whereas skin genes most often correlated with DLCO. Abbreviations: PBMC, peripheral blood mononuclear cells; DLCO, diffusion capacity of the lungs for carbon monoxide; MRSS, modified Rodnan skin score; FVC, forced vital capacity.

| **Tissue** | **Severity** | **Negative** | **Positive** |
| --- | --- | --- | --- |
| PBMC | DLCO | 141 | 79 |
|  | Forearm skin score | 13 | 15 |
|  | MRSS | 129 | 226 |
| Skin | DLCO | 688 | 553 |
|  | FVC | 427 | 280 |
|  | MRSS | 340 | 305 |

For PBMCs, there were significant correlations between gene expression and DLCO (**Table ST8**), forearm skin score (**Table ST11**), and MRSS (**Table ST13**). The genes negatively correlated with DLCO were enriched for type 1 interferon signaling, as well as interferon responsive factor and STAT2 transcription factors (**Table ST9**). Proteasomal antigen processing and presentation pathways (REAC:R-HSA-1236975, GO:0002479, GO:0002474, GO:0042590) were enriched primarily due to negative correlations of *HLA-B*, *PSMA4*, *PSMC2*, *PSME1*, *PSME2*, and *TAPBP* with DLCO. Positive correlations with DLCO included signal peptide CUB domain and EGF-like domain-containing 3 (*SCUBE3*; 0.74 correlation [corr]) and antisense transcript *AC008440.1* (0.63 corr). The PBMC genes that positively correlated with DLCO were only enriched for targets of microRNA hsa-miR-6082 (*DTWD2*, *FXN*, and *ZFP30*; **Table ST10**).

Forearm-specific skin score correlations (**Table ST11**) were more difficult to interpret. The negative correlations did not show any pathway enrichments. The phosphoribosylformylglycinamidine synthase activity pathway was enriched, but it was due to a single gene (*PFAS*; **Table ST12**).

There were also correlations between PBMC gene expression and MRSS (**Table ST13**). Genes negatively correlated with MRSS were enriched for pathways associated with protein folding, unfolded proteins, and ER stress (**Table ST14**). There were a few pathways associated with positively correlated genes, including some related to mitochondria (**Table ST15**).

One might assume that the differentially expressed genes in case/control gene expression studies may also be the key genes and pathways that drive progression and therefore may correlate with disease severity. This does not appear to be the case for SSc PBMCs, as most of the genes with significant correlations to severity were not detected in the case/control analysis (**Fig. 6**).

**Fig. 6.**
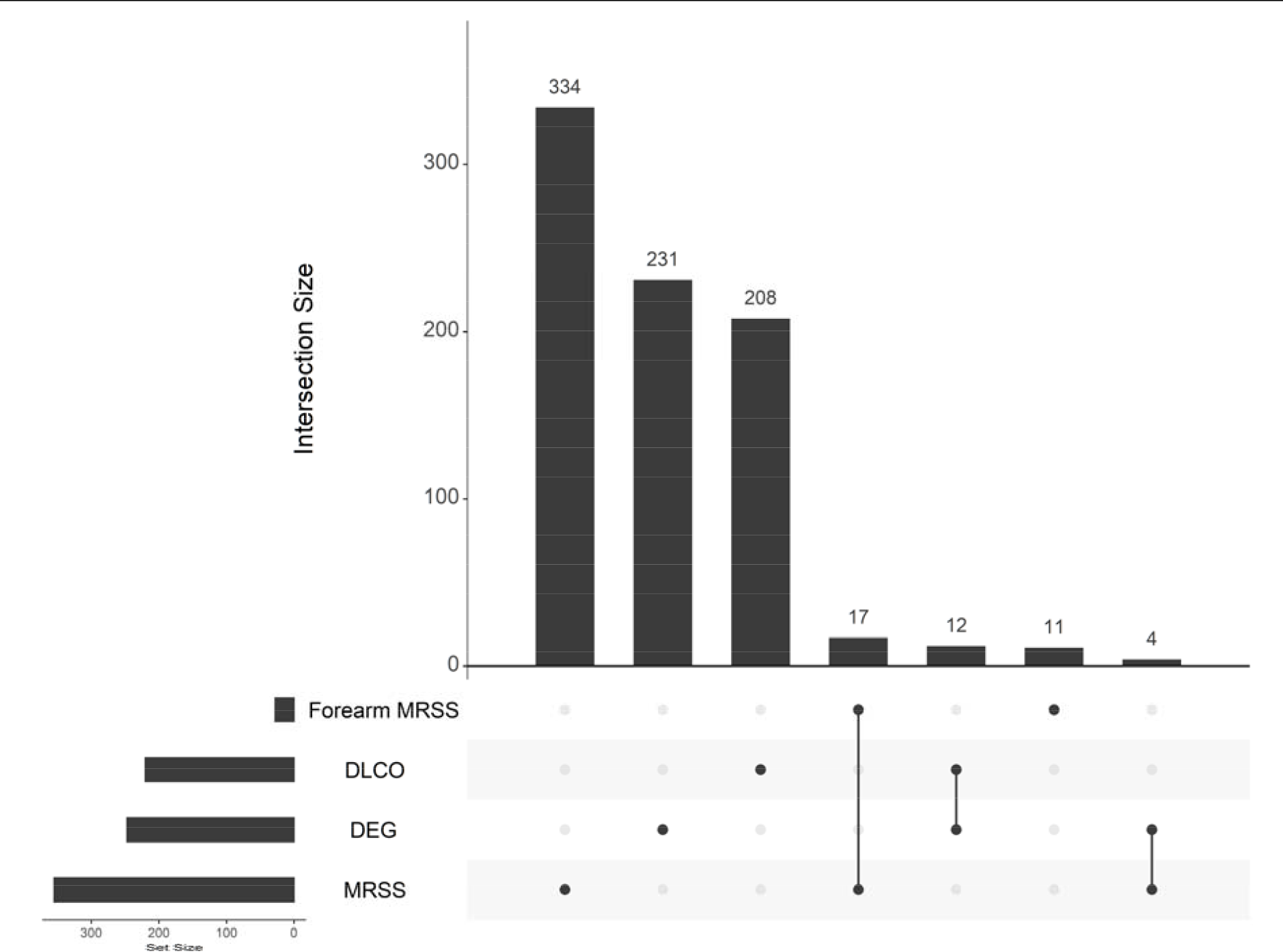
PBMC differentially expressed genes do not explain clinical trait correlations. SSc PBMC transcriptomes had correlations with Forearm MRSS, DLCO, and MRSS by weighted gene co-expression analysis. The data for each category are plotted in an UpSet. The bar plots to the left indicate the total number in the given category. The upper bar plots and corresponding connected dots give the number for the intersection. Overall, only 12 genes were both differentially expressed and trait correlated for PBMCs. This highlights the unique information available from each method. Abbreviations: DEG, differentially expressed genes; DLCO, diffusion capacity of the lungs for carbon monoxide; MRSS, modified Rodnan skin score.

The SSc skin transcriptomes had significant correlations with DLCO (**Table ST16**), FVC (**Table ST19**), and MRSS (**ST21**). For DLCO, many negative correlations were with ribosomal proteins, leading to enrichment of pathways related to ribosome function (**Table ST17**). SSc skin genes that positively correlated with DLCO did not fall into a consistent theme (**Table ST18**). Negative correlations with FVC were enriched for sterol/cholesterol biosynthesis and alpha-linolenic/linolenic acid metabolism pathways (**Table ST20**). The positive correlations with FVC did not have any significant pathway enrichment.

Skin genes negatively correlated with MRSS were enriched in cell fate and synaptic categories (**Table ST22**). Genes positively correlated with MRSS were enriched for genes associated with the extracellular matrix (**Table ST23**). This is a nice confirmation, as increasing MRSS should indicate increasing fibrosis, secondary to increased matrix deposition. The major connective tissue themes were extracellular matrix and collagen fibers. There was a single enrichment category for TGF β binding (GO:0050431) due to positive correlations of MRSS with *LRRC32*, *TGFBR2*, and *TWSG1*. LRRC32 (also known as GARP) is expressed in activated regulatory T-cells, binds to TGF-β 1, and is required for the surface expression of latent TGF-β 1 in these cells (Tran et al., 2009). This interaction is important for the immunosuppressive function of regulatory T-cells, as monoclonal antibodies targeting GARP/TGF-β1 complexes reduce the immunomodulatory effect of these cells (Cuende et al., 2015).

Similar to what we observed in PBMCs, there is little overlap between the genes differentially expressed in SSc skin and those that correlated with lung function or skin fibrosis parameters (**Fig. 7**). Overall, 90% of the differentially expressed genes did not correlate with any clinical parameters, and 93% of the clinical trait-associated genes were not differentially expressed. This highlights the disparity between the two methods and suggests that novel targets for clinical treatment and biomarkers may be identified using severity correlation rather than case/control differential expression.

**Fig. 7.**
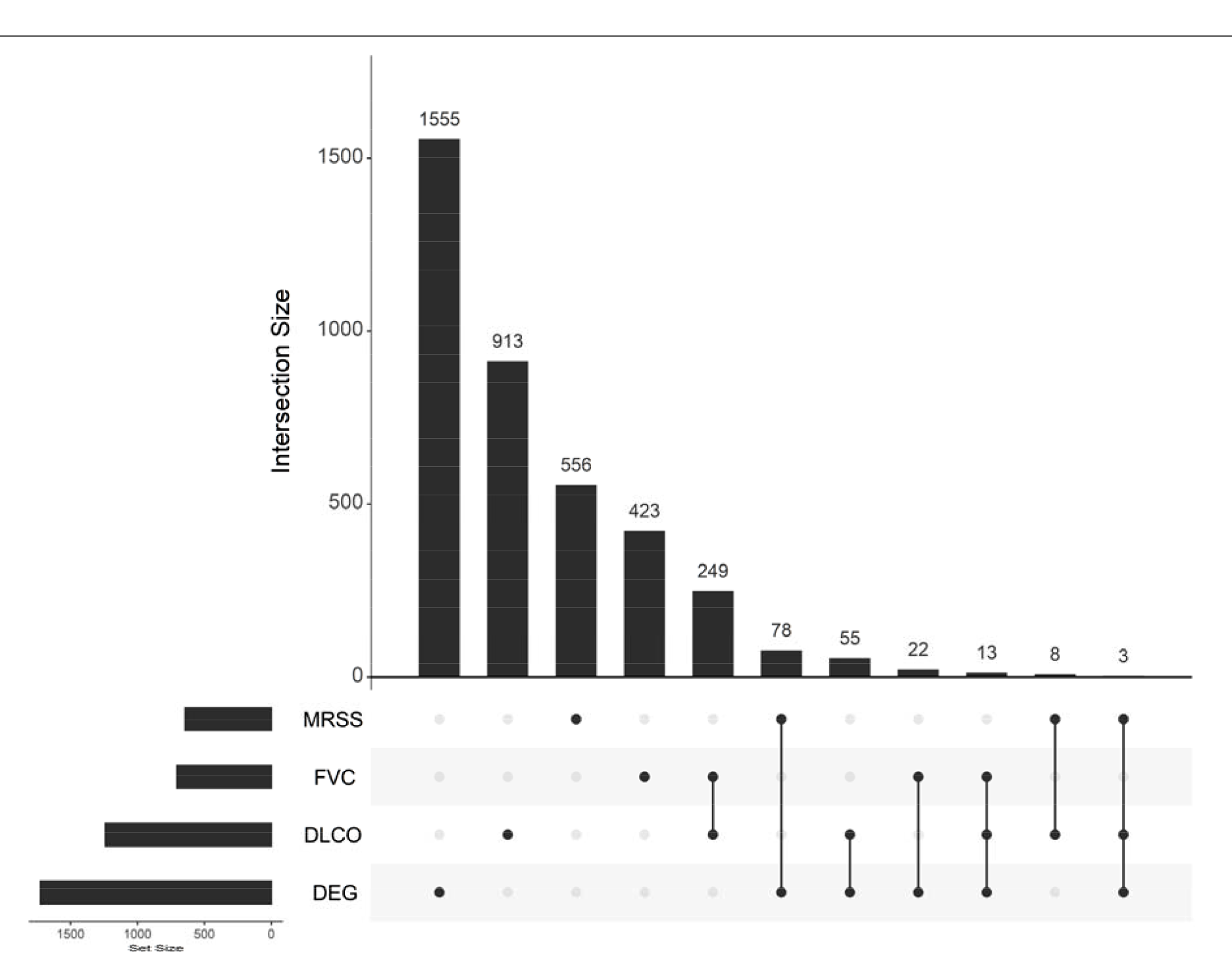
Genes in SSc skin that correlate with clinical severity are usually not differentially expressed Skin genes significantly correlated with skin fibrosis (MRSS) and lung function parameters (DLCO and FVC). There were many more correlated genes and differentially expressed genes in the skin,as might be expected since it is an affected tissue. 171 genes were both differentially expressed in the case/control analysis and significantly correlated with at least one trait. Most of the significant correlations were with total MRSS and DLCO. However, the general trend was that a gene would either be detected as differentially expressed or trait correlated rather than overlapping.

Some genes were correlated with the same trait in both skin and PBMCs (**Table ST24**). For each gene, we considered the tissues concordant if the direction of effect was the same in each tissue, and discordant if the direction of effect was opposite. There were only overlaps between tissues for DLCO and MRSS. DLCO had 12 concordant genes and 7 discordant genes between tissues. MRSS had 11 concordant and 2 discordant genes. These two gene sets are intriguing to consider for further study as molecular biomarkers of disease activity, regardless of whether the effect is concordant or discordant, as a blood draw might be as informative as a skin punch biopsy.

### Non-coding RNAs SOX9-AS1 and ROCR are central, highly connected in the SSc skin gene-gene **correlation network**

Given that we had a list of genes that correlated with different traits and their normalized expression, the next thing we looked at was the gene-gene correlation network. We focused only on genes that significantly correlated with at least one clinical trait. Examining the network characteristics can help identify genes that act as signaling hubs or are otherwise have co-expression with other members of the network.

After ranking each gene by degree and page rank (measures of network connectedness and centrality), the top-ranked PBMC gene was 2’-5’-oligoadenylate synthetase 2 (*OAS2*) with a degree of 29 and page rank of 0.017 (**Table ST25**; **Fig. 8**). The oligoadenylate synthetases are interferon-response genes, which is consistent with the increased type 1 interferon signaling signatures in SSc PBMCs. The 4^th^ highest ranked gene, *EPSTI1*, is linked to macrophage function. It’s thought to have a key role in classical M1 polarization of macrophages, as a murine knockout of *Epsti1* has few M1 polarized macrophages along with a significant expansion of M2 polarized macrophages (Kim et al., 2018). The top-ranked genes in SSc PBMCs (all genes with a degree of at least 10) are inflammatory genes that correlated with DLCO.

**Fig. 8.**
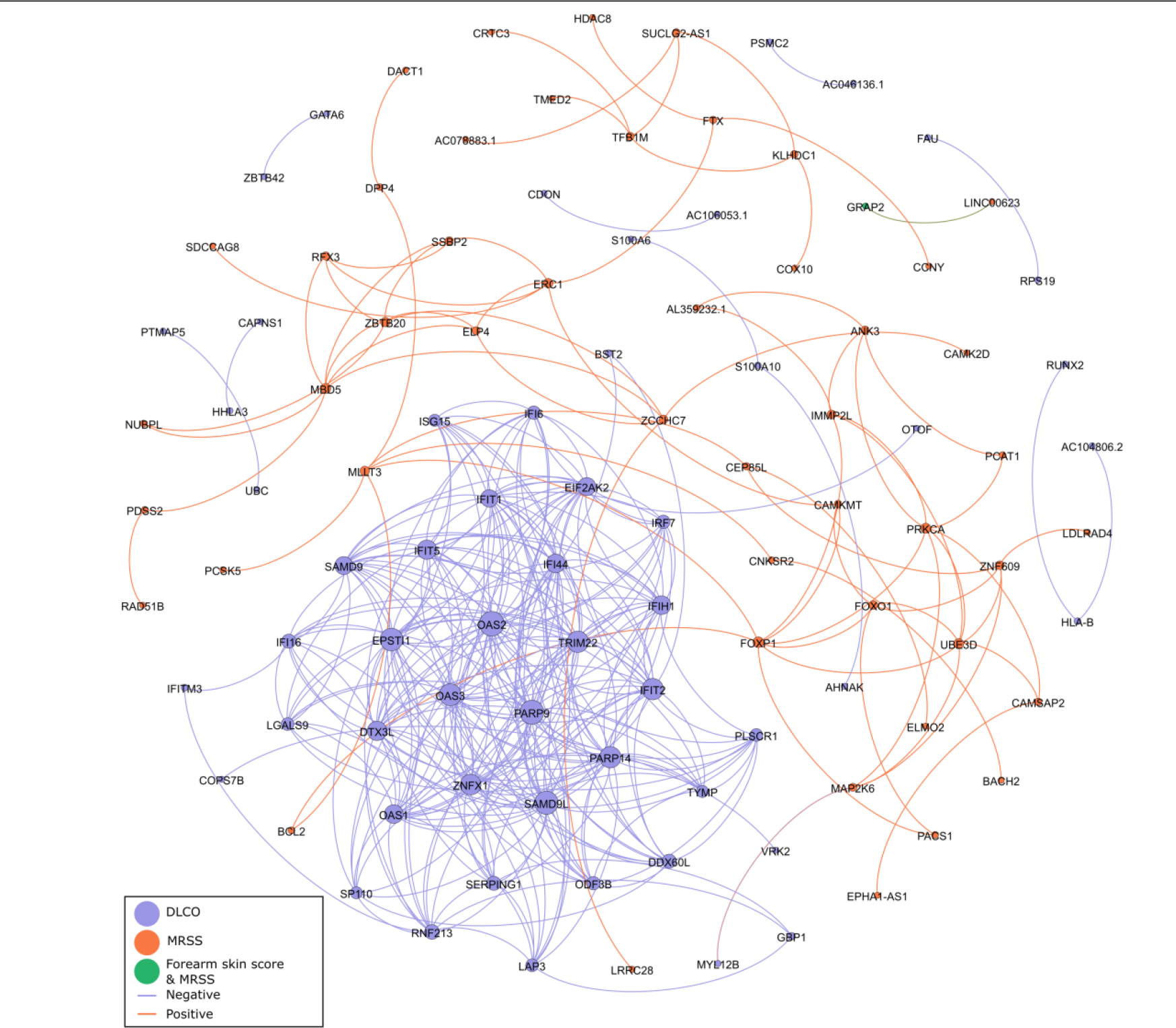
Interferon-responsive genes are the most highly connected trait-correlated genes in SSc PBMCs. Shown is a network diagram for genes correlated with at least one clinical trait. Each node is an individual gene, sized by weighted degree and filled by the trait association. Each edge is colored for whether the gene-gene correlation is positive or negative, with a minimum cutoff of 0.80. The interferon-responsive genes, such as *OAS1*, *OAS2*, and *OAS3*, and *IFIT1/2*, were the most highly interconnected for PBMCs. Abbreviations: PBMC, peripheral blood mononuclear cells; DLCO, diffusion capacity of the lungs for carbon monoxide; MRSS, modified Rodnan skin score.

Conversely, the top 41 genes in the SSc skin network (ranked by degree and page rank) all correlated with MRSS (**Table ST26**; **Fig. 9**). The top-ranked SSc skin gene was an antisense transcript, *SOX9-AS1* (degree = 49; page rank = 0.005). This highlights an important advantage of using stranded RNA library kits: with an unstranded kit it is impossible to tell sense from antisense transcripts at the same locus. The protein-coding *SOX9* transcript was correlated with MRSS as well but had a lower degree of 9. One proposed role for *SOX9-AS1* is as a microRNA sponge, i.e. the antisense transcript competes with *SOX9* as a target for repression. Therefore, increased expression of *SOX9-AS1* could lead to increased *SOX9*. Consistent with this idea, knocking down *SOX9-AS1* in the human Huh7 hepatocellular carcinoma cell line decreased the expression of *SOX9* (Zhang W. et al., 2019). The same study also showed that *SOX9-AS1* is a target of the SOX9 transcription factor and that treatment with Wnt/ β-catenin agonists canceled the effect of the *SOX9-AS1* knockdown. The long non-coding RNA *ROCR* is also a highly connected gene in the SSc skin network (degree = 44; page rank 0.004) and may have a direct role in fibrosis. It is located in the same genomic locus as *SOX9* and *SOX9-AS1*. Knocking down *ROCR* during chondrogenic differentiation of human mesenchymal stem cells results in lower production of matrix genes *COL2A1* and *ACAN* (Barter et al., 2017). *ROCR* may also be a target of SOX9 since overexpression of *SOX9* recovers the chondrogenic differentiation program in *ROCR* knockdown cells. It is worth noting that *SPAG17* expression in SSc skin was negatively correlated with MRSS (-0.51 correlation). Since *SPAG17* is a low expression transcript, we were unable to generate a *SPAG17* network to identify co-expressed genes. Therefore further study is required to understand the genes co-expressed with *SPAG17* in relevant skin cell types, such as fibroblasts.

**Fig. 9.**
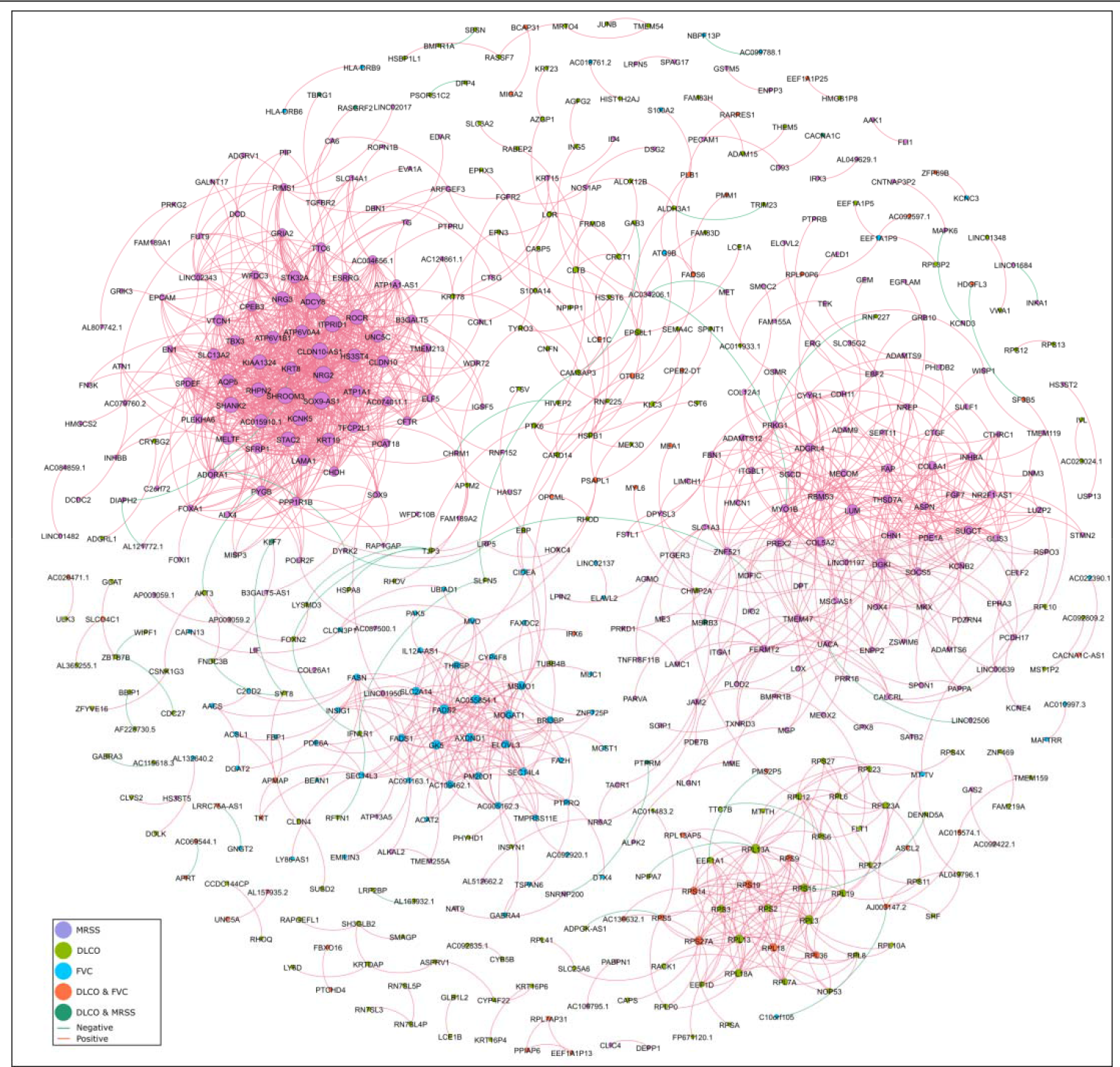
*SOX9* locus genes are among the most highly connected trait-associated genes in SSc skin Shown is a network diagram for genes correlated with at least one clinical trait in the skin. Each node is an individual gene, sized by weighted degree and colored by the trait association. The edges between nodes are colored for whether the gene-gene correlation is positive or negative, with a minimum cutoff of 0.80. Some genes correlated with fibrosis formed a relatively separate sub-network from those associated with lung function or both DLCO and fibrosis. The most connected genes in this sub-network included *SOX9-AS1* and *ROCR*, which are both non-coding and located at the *SOX9* locus. The overlap of correlations between fibrosis and lung function were primarily ribosomal proteins. A separate sub-network of MRSS-correlated genes was enriched for matrix proteins, such as *COL5A2* and *COL8A1*. Abbreviations: DLCO, diffusion capacity of the lungs for carbon monoxide; FVC, forced vital capacity; MRSS, modified Rodnan skin score.

## DISCUSSION

Systemic sclerosis is a complex and progressive inflammatory/fibrotic disease. In the current study, we show that the sensitivity of RNA-Seq can lead to new discoveries and that applying complementary approaches to the same data can reveal distinct trends. There are advantages to using strand-specific, ribosomal depletion RNA-Seq library kits: increased ability to detect non-polyadenylated transcripts, increased sensitivity for nascent and short half-life transcripts, and the ability to distinguish overlapping antisense transcripts. However, a disadvantage is a reduced power for transcript-level discoveries. For our data, focusing only on spliced transcripts would have used only a fraction of the available sequencing data, since much of the aligned sequence mapped to introns. There is also distinct value to using different tissues. Blood samples are informative in their own right, but comparing skin punches between SSc patients and controls reveals different gene sets more directly related to the ongoing molecular pathology.

The predominant signal in PBMCs was for type 1 interferon signaling, with enrichment for targets of IRF5 and IRF9. M1 polarized macrophages are known to have high *IRF5* expression (Krausgruber et al., 2011). We observed that *CD86*, another marker of M1 macrophages, was increased in SSc PBMCs. Previous work has shown that a mix of M1 and M2 macrophages in the blood is associated with interstitial lung disease in systemic sclerosis (Trombetta et al., 2018). The lack of an increase of M2 markers doesn’t necessarily exclude their presence. Bulk RNA-Seq is not ideal for identifying cell populations and the question of macrophage fraction (and activation state) in SSc PBMCs would be better answered with single-cell RNA-Seq, flow cytometry, or mass cytometry.

The skin data provide an even more informative perspective. The most significantly different gene in the entire analysis was *SPAG17*, which has not traditionally been a top dysregulated gene in SSc transcriptome studies. Genes are often named for the tissue in which they are first discovered, which may or may not be the only tissue they’re expressed in or even the tissue with the highest expression.

SPAG17 is a central pair protein that is critical in the formation of primary cilia, and its knockout is neonatal lethal in mice (Teves et al., 2013). As suggested by the name, alterations in *SPAG17* can lead to infertility in mouse models and for humans with certain rare missense variants (Kazarian et al., 2018, Xu et al., 2018). But it is important to bear in mind that this gene has critical functions beyond sperm motility since it is part of the primary cilia. Missense changes in *SPAG17* can cause abnormal bone length (Cordova-Fletes et al., 2018, Teves et al., 2015). Common variants in *SPAG17* are associated with body length in early life and height in adulthood (Kim et al., 2010, van der Valk et al., 2015).

Novel mutations and rare variants in components of the primary cilia can lead to primary ciliary dyskinesia (**PCD**). The most common effects are in the ears (chronic ear infections, hearing loss), sinuses (chronic sinus congestion), and lungs (recurrent pneumonia, chronic cough). *SPAG17* rare variants can lead to a PCD-like phenotype in mice and have been shown to cause human PCD as well (Abdelhamed et al., 2020, Andjelkovic et al., 2018). The PCD phenotypes are largely driven by the altered ability of the cilia to beat. In systemic sclerosis skin, a defect of beating cilia doesn’t make the most sense. Non-motile primary cilia are involved in signaling. One potential hypothesis is that primary cilia have an anti-fibrotic signaling role. The reduced expression could then lead to increased profibrotic signaling. Yet to be elucidated is whether mouse *Spag17* hypomorphs have increased fibrosis susceptibility, which cells in the skin are affected by reduced *SPAG17* expression, and why the *SPAG17* expression is decreased in the first place, i.e. whether it is a primary or secondary effect. One possibility is that *SPAG17* is lost in the transition from fibroblast to myofibroblast. Further research will be required to dissect these possibilities, particularly since the mouse germline knockout is neonatal lethal.

We detected a decreased expression of *LGR5*in SSc skin. The recent finding that *LGR5* is a marker for mouse intestinal villi tip telocytes begs the question of whether it is a marker of skin telocytes. The intestinal tip telocytes had increased expression of *Wnt5a* compared to the intestinal crypt telocytes, suggesting that non-canonical Wnt signaling is important in those cells (Bahar Halpern et al., 2020). Skin telocytes are known to form multiple contacts to ECM and other cells, such as adipose cells and fibroblasts (Rusu et al., 2012). One possible function of these interconnections is to provide support for the other cells within the skin matrix. It is, however, also possible that these cells are critical for the transduction of signals within the skin, and therefore may play a direct role in the evolution of fibrosis in SSc.

SSc skin had enrichment for pathways related to classical (antibody-mediated) complement activation. There is a growing body of evidence that complement activation and subsequent damage play a role in systemic sclerosis endothelial damage. The terminal effector of complement damage, the membrane attack complex, has been observed in the small vessels of affected systemic sclerosis skin and the muscle endothelium of patients with systemic sclerosis-associated myositis (Corallo et al., 2017, Scambi et al., 2015). This indicates local tissue damage, regardless of how it is triggered, may be complement-mediated. This possibility is perhaps even more evident in scleroderma renal crisis (**SRC**), which has a sudden onset of severe hypertension and acute renal failure. The kidneys of some individuals with SRC show deposition of complement C3b in renal arterioles (Okroj et al., 2016, Perez et al., 2019). Complement deposition without substantial inflammation and the presence of thrombotic microangiopathy is also a hallmark of atypical hemolytic uremic syndrome (**aHUS**). Familial aHUS is often caused by aberrant regulation of complement activation, particularly via genetic variants in complement factor H (Noris et al., 2010). The first-line therapy for aHUS is eculizumab, a monoclonal antibody to C5 (Cofiell et al., 2015). There is some evidence that eculizumab might also be effective for SRC (Devresse et al., 2016, Uriarte et al., 2018). However, this still leaves unanswered whether endothelial complement activation is a major driver of SSc skin vasculopathy.

The network analysis of PBMC and skin transcriptomes with skin fibrosis and lung function parameters demonstrated that both tissues are informative for different traits, opening the possibility of the development of minimally invasive, quantitative biomarkers of disease activity. This would be a boon to clinical trials, particularly if observable parameters, such as MRSS, don’t tell the whole story about internal disease progression. A particularly interesting facet of the clinical trait correlation in our study is the suggestion that SOX9 is a critical player in fibrosis. Expression of the *ROCR* and *SOX9-AS1* non-coding RNAs, as well as SOX9 itself, was significantly correlated with fibrosis in SSc skin.

Importantly, *SOX9-AS1* and *ROCR* were two of the most highly interconnected genes in the SSc skin gene-gene co-expression network, suggesting a key role in mediating the progression of fibrosis. Both β-catenin signaling with catenin (Zhang W. et al., 2019). We previously demonstrated that blocking Wnt/β C-82 restored subdermal adipogenesis in patients with SSc (Lafyatis et al., 2017). However, since these are non-coding RNAs, it is currently unclear whether their effect is primarily through a role in regulating *SOX9* or through an alternative mechanism unrelated to *SOX9*. In particular, non-coding RNAs also act as linkers between DNA and protein. One such example is the *HOTAIR* non-coding RNA that mediates repression of some *HOX* loci by recruiting complexes to repress chromatin in those regions (Rinn et al., 2007, Tsai et al., 2010). Further dissection of the role and mechanism of action of *SPAG17* in profibrotic signaling and a better understanding of the roles of *ROCR* and *HOX9-AS1* in the skin are intriguing areas for future research.

## MATERIALS & METHODS

### Clinical assessment

We completed standardized evaluations to establish SSc diagnosis, as well as presence/severity of organ involvement, as previously described (Hinchcliff et al., 2013, Johnson et al., 2015). We determined modified Rodnan Skin Score and local (forearm) skin scores at each visit. The high-resolution CT of the chest, echocardiography, and pulmonary function testing was performed as standard of care (Richardson et al., 2016).

### Sample collection and storage

At each visit, we collected peripheral blood mononuclear cells (**PBMCs**), serum, and skin punch biopsies in addition to the clinical data. We collected a single sample set from controls. When possible, we collected follow-up samples at six months from systemic sclerosis patients. For PBMC isolation, we collected 8 mL of blood in CPT tubes. We separated PBMCs from red cells using Ficoll density gradient centrifugation, spinning the tubes at 1500-1800 rcf with no brake for 20-30 minutes at room temperature (**RT**). We discarded the supernatant, gently vortexed the pellet to resuspend, and added 10 mL of PBS to wash the cells. We centrifuged at 300 rcf for 10 minutes without braking and removed the supernatant. We lysed the cells by adding 700 µL of Qiazol followed by vortexing. We stored the lysed slurry at -80°C until RNA extraction. For serum isolation, we collected 10 mL of blood in SST tubes and stored the serum in 500 µL aliquots at -80°C for future use.

We obtained two separate 3 mm skin punches per individual per visit. We collected one biopsy in formalin, processed it for routine histology, and stored it in a paraffin-embedded tissue block for future immunohistochemistry. We treated the other biopsy overnight with RNAlater at 4°C to stabilize the tissue RNA. We stored the stabilized punches at -80°C.

### RNA extraction

We allowed RNAlater treated samples to thaw at 4°C on ice. Once thawed, we removed the RNAlater solution and minced the skin punches into smaller pieces. We then disrupted and homogenized the samples using a Qiagen Tissueruptor II (#9002755). We extracted total RNA from the homogenized samples using the miRNeasy Minikit (#217004) according to the manufacturer’s instructions. We stored the RNA at -80°C until further use.

### RNA-Seq library preparation

We used the Clontech SMARTer stranded total RNA high input sample preparation method to generate the RNA-Seq libraries (#634876). In brief, we removed ribosomal RNA (**rRNA**) by incubating the RNA with DNA probes that target rRNA, followed by treatment with RNase H to selectively degrade the RNA component of RNA-DNA hybrids. We then removed DNA probes by treatment with DNase I. We used the rRNA-depleted samples for cDNA generation and PCR amplification. The resulting libraries represent total RNA, and read 1 is in sense orientation to the initial transcript. We used unique i5 / i7 combinatorial TruSeq index pairs for each member of a sample pool. We then sequenced the pools in paired-end mode on either a HiSeq2500 (101 bp reads) or HiSeq3000 (150 bp reads) sequencer at the Genome Technology Access Center at Washington University in St. Louis.

### Sequence generation, pre-processing, & differential expression analysis

For pre-processing, we removed adapters and trimmed low-quality 3’ sequences with cutadapt v1.17 (Martin, 2011). For reference-based alignment, we aligned with RNA-STAR v2.6.0c using GRCh38 release 85 of the human genome plus contigs for several human pathogens and common cell culture contaminants (details in supplement). We counted reads overlapping genes in two different ways using featureCounts v1.5.1 (Dobin et al., 2013, Liao et al., 2014). For mature transcripts, we counted against the Gene Transfer Format file of known genes for GRCh38. This would include reads fully contained within exons and those that crossed exon-exon boundaries. For both unspliced and mature spliced transcripts, we counted all reads overlapping a known gene (between transcript start and end) on the correct strand. Some libraries were sequenced in multiple lanes. We summed the read counts per gene across all runs for a given sample library before calculating differential expression. We used DESeq2 (v1.24.0) to calculate differential expression (Wald test) between SSc and control for both tissues, and normalized expression (variance stabilizing transform and regularized logarithm methods) in R v3.5.1 or v3.6.0 (Love et al., 2014). We used R for miscellaneous statistical calculations and figure generation as well.

### Weighted gene co-expression network analysis with disease severity

We used Weighted Gene Co-expression Network Analysis (**WGCNA**) to correlate expression with MRSS, forearm skin score, FVC, and DLCO (Langfelder and Horvath, 2008). The input expression data was the total read counts for each gene adjusted for batch effects by ComBat-Seq (Zhang et al., 2020) and normalized in DESeq2 using the vst method. We included initial and follow-up samples for any SSc subtype for which clinical data were available. In WGCNA, we tested soft-thresholding powers from 1 – 30, with a minimum module size of 10 and a scale-free topology soft threshold of 0.85.

### Gene-gene correlation network construction and network statistics

We calculated the pair-wise correlation between all genes using the WGCNA package biweight midcorrelation with 0.05 maximum fraction of outliers. For these networks, we only included genes that were significantly correlated with at least one clinical trait. We retained edges between genes if the absolute correlation was at least 0.80. We imported the pair-wise correlations (edges) and gene-trait association information (nodes) into Gephi (v0.9.2). We calculated all network statistics using the Network Overview options. For visualization, we colored nodes based on the clinical traits they were associated with. Node size was based on ranked weighted degree, with a minimum size of 10 and a maximum size of 45. Edges were colored based on whether the correlation was positive or negative. The network was arranged using the Fruchterman Reingold algorithm with an area of 20,000 and a gravity of 5.0. The network was exported as an SVG, and the final key was generated using InkScape (v0.92.4).

### Pathway enrichment

We calculated pathway enrichments using gProfileR (ve96_eg43_p13_563554d) with the following options: return only significant adjusted enrichments, unordered query, no electronic Gene Ontology annotations, Best Parent (moderate) filtering, maximum category size 500, and to adjust the significance threshold using the g:SCS (cumulative hypergeometric probability) method (Raudvere et al., 2019). For differentially expressed genes, we tested increased and decreased genes independently. For trait associations, we tested significant positive and negative correlations separately.

## DATA AVAILABILITY

Data used for analysis, such as gene counts used for DESeq2 and demographic information per sample, are available from FigShare. https://figshare.com/projects/2021_Roberson_lab_systemic_sclerosis_transcriptome_data/118698

Code used for data analysis is available as a repository on GitHub. https://github.com/RobersonLab/2021_ssc_rnaseq

These were prospectively collected samples for controlled data access. We are currently in the process of depositing the FASTQ files into dbGAP.

## ACKNOWLEDGMENTS

This study was supported by the Rheumatic Disease Core Center (NIAMS P30-AR048335; EDOR, LC, JPA) and the Rheumatic Diseases Research Resource-based Center (NIAMS P30-AR073752; EDOR) at Washington University. JV was supported by the Northwestern University Clinical and Translational Sciences Institute (UL1-TR000150). JPA and EDOR were supported by the Washington University in St. Louis Institute of Clinical and Translational Sciences [ICTS] (UL1-TR000448). DJM-H was supported by an NIH training grant (T32-AR007279-36). Some of the analysis was performed with support from the Washington University Center for High-Performance Computing (S10-OD018091). Sequencing data were generated at the Genome Technology Access Center (GTAC@MGI) at Washington University School. GTAC is partially supported by the Siteman Cancer Center (P30- CA91842) and the ICTS (UL1-TR000448). The research presented herein represents the views of the authors and does not necessarily reflect the views of the NIH.

