## Supplemental Text & Methods for "RNA-Seq analysis identifies novel roles for the primary cilia gene *SPAG17* and the *SOX9* locus non-coding RNAs in systemic sclerosis"

### SUPPLEMENTAL MATERIAL

#### Supplemental data

***VEDOSS shows blood and skin transcriptional profiles intermediate between control and SSc, while systemic sclerosis sine scleroderma skin transcriptomes are indistinguishable from lcSSc and dcSSc***

We were able to identify genes differentially expressed in SSc skin and PBMCs, but this analysis doesn't address how variable these signatures are amongst individuals. We, therefore, chose to perform dimensionality reduction to address this question. We included initial and follow-up samples for control, limited cutaneous systemic sclerosis, diffuse cutaneous systemic sclerosis, systemic sclerosis sine scleroderma, and very early diagnosis of systemic sclerosis. We used the normalized expression as input for principal components analysis (separately for each tissue), limiting the genes to the top 100 most significant as ranked by adjusted p-value. For PBMCs, the effect seems to correlate with the extent of disease (**Fig. S1**). The SSS, VEDOSS, and some limited cutaneous systemic sclerosis samples cluster among the controls. Some of the lcSSc samples, and all of the dcSSc samples, separate clearly from the controls. This suggests that the level of immunological activation in PBMCs may correlate with the extent of skin involvement. It doesn't, however, indicate whether the enhanced activation is a phenomenon secondary to the increased connective tissue involvement, or if the increased connective tissue involvement is caused by the increased immune activation.

We used the same approach with the normalized skin expression data, but with different results. In the skin, all SSc samples separate clearly from the controls (**Fig. S2**). The SSc samples that are closest to the controls are the VEDOSS sample and one limited cutaneous sample. It's therefore likely this is a true VEDOSS sample, as it isn't exactly control-like, but doesn't yet have a clear SSc transcriptome-wide signature. Unexpectedly, the systemic sclerosis sine scleroderma sample, which would not have extensive skin fibrosis, clusters along with the limited and diffuse cutaneous systemic sclerosis samples. These data, therefore, suggest that there may be substantial transcriptomic disruptions in SSS that are do not lead to fibrosis but are nonetheless important to disease progression and classification.

#### Supplemental methods

***Sequencing pool generation***

We uniquely identified each member of a sequencing library pool by using a distinct combination of forward and reverse indexes (i5/i7 combinatorial indexing), i.e. the forward and reverse primer pair was unique per sample. We determined the concentration of each library using the Qubit high-sensitivity DNA assay (Q32854; ThermoFisher Scientific, Waltham, MA). We confirmed the integrity of each library by combining 1 µL of each library in a pool, adjusting the final concentration 10 nM, and sequencing approximately 500,000 – 1,000,000 reads on an Illumina MiSeq by spiking the pool in with another sample set. We used the calculated reads / µL, which would intrinsically account for differences in library insert size distribution, to determine the volume of each library to combine for the final sequencing pool.

***Sequence generation and raw data pre-processing***

We generated sequencing data using both the Illumina HiSeq2500 and the HiSeq3000. The HiSeq2500 runs were paired-end 101 bp cycle read length, and the HiSeq3000 runs were paired-end 150 bp cycle read length. The HiSeq3000 runs required a minimum of 10% PhiX spike-in to avoid QC failure during the cluster identification step. The Clontech SMARTer libraries use the Moloney Murine Leukemia Virus Reverse Transcriptase with template switching to generate cDNA. This results in the non-templated addition of a triple C base at the 3' end of the first strand. The low complexity in the first few reads triggers a QC fail on the HiSeq3000 without the addition of PhiX (though any sufficiently complex library would suffice). We de-multiplexed the raw FASTQ files using the expected unique combinatorial indexes. We cleaned the raw FASTQ files by removing contaminating sequencing adapter and trimming low-quality bases from the 3' end of reads with cutadapt (v1.17; Python v3.5.5).

Cutadapt options:

--cut 6, --max-n 0.20, -q 10, -m 30, -O 5, -a AGATCGGAAGAGC, -A SSSAGATCGGAAGAGC

We cut 6 bases from the 5' end of read 1 because of the low complexity of the first few bases, which was confirmed by FastQC. The degenerate S base for the read 2 adapter accounts for the non-templated MMLV base addition, allowing for any GC read in the first 3 bases to be removed as well.

***Reference-based alignment***

We used the human genome version GRCh38 release 85 top-level chromosomes (soft-masked) from Ensembl for our reference sequence. The soft masking represents repeats as lower case letters rather than N bases. In addition, we include some additional sequences to speed the alignment step. This includes some common internal spike-in standard controls from Ambion (#AM1780; ThermoFisher Scientific, Waltham, MA), the PhiX phage sequence Phi X 174 (genbank #9626372), Moloney Murine Leukemia Virus (#NC_001501.1), common cell culture contaminants (*Acholeplasma laidlawii pg8a*, #161984995; *Mycoplasma fermentans m64*, #318037813; *Mycoplasma hominis*, #269114774; *Mycoplasma hyorhinis MCLD*, #330723203), and some human pathogens (chikungunya virus, #615794504; cytomegalovirus, #559798448; Epstein-Barr virus, #557368242; hepatitis B, #666689229; hepatitis C, #669687798; human immunodeficiency virus 1, #9629357; human papilloma virus, #604724199).

We used 2-pass mode for RNA-STAR v2.6.0c. The first pass was to identify splice junctions in all samples. The Nprocessors for the number of threads varied depending on what resources were available.

STAR 1st pass options:

--runMode alignReads --runThreadN Nprocessors --readFilesCommand zcat --outFilterMultipmapNmax 10 --outFilterScoreMinOverLread 0.45 --outFilterMatchNminOverLread 0.45 --outSAMtype BAM Unsorted outReadsUnmapped Fastx

We extracted all novel junctions detected in any sample and kept novel any with at least 10 reads in the overall set. We then ran the second-pass alignment including these novel junctions (inserted "on-the-fly") to get a final alignment.

STAR 2nd pass options:

--runThreadN Nprocessors --readFilesCommand zcat --outFilterMultimapNmax 10 --sjdbFileChrStartEnd OurNovelJxnFile.txt --outFilterScoreMinOverLread 0.45 --outFilterMatchNminOverLread 0.45 --outSAMtype BAM Unsorted outReadsUnmapped Fastx

***Counting reads with featureCounts***

We used the unsorted BAMs to determine the number of reads consistent with each gene in the Ensembl GTF for our genome build using featureCounts (v1.5.1).

Gene-based counting options:

--donotsort -p -a /path/to/gene_model.gtf -Q 10 -T Nprocessors -s 1 -o output.count

Since we used a ribosomal depletion library method, we had a substantial number of reads that contained intron sequences. Instead of using the GTF to guide this step, we used stranded counting on the region of each gene, starting at the gene start and ending at the gene end.

Total gene read counting options:

--donotsort -p -a /path/to/gene_chrom_start_end_file.txt -F SAF -Q 10 -T Nprocessors -s 1 -o all_gene_reads.count

***Library metrics***

For each STAR alignment, we had output for the number of reads mapping to different parts of the genome, categorized as "unique", "multiple", "too many", and "unmapped". Unique mapping reads are perhaps better for estimating transcript abundance than the others. High fractions of unmapped reads suggest lower input sample quality or issues in library preparation. We also determined the insert size distribution and RNA-Seq metrics using Picard tools v2.7.1 (CollectInsertSizeMetrics, CollectRnaSeqMetrics). We used the Ensembl GRCh38 release 85 locations of nuclear ribosomal RNA, mitochondrial ribosomal RNA, and tRNAs to supply “ribosomal” locations to Picard tools.

### Supplemental table descriptions

Differential expression results: ST1, ST4

gene_id: Ensembl gene id.

symbol: associated gene symbol.

gene_biotype: Ensembl gene biotype annotation.

FoldChange: case / control fold-change measurement from DESeq2.

pval: Wald test p-value from DESeq2.

qval: False-discovery rate corrected p-value.

baseMean: base mean value from DESeq2.

log2FoldChange: log2 fold-change from DESeq2.

lfcSE: log fold-change standard error.

stat: Wald statistic from DESeq2.

gProfileR pathway results: ST2-3, ST5-6, ST9-10, ST12, ST14-15, ST17-18, ST20, ST22-23

adjusted_p_value: gProfileR g:SCS corrected p-value.

source: Data source. Example – GO:CC = Gene Ontology, Cellular Compartment. REAC = Reactome.

term_id: known pathway term ID.

term_name: known pathway name.

symbols: list of gene symbols in this pathway.

Intersections: list of Ensembl gene IDs in the intersection.

Differential expression overlap: ST7

gene_id: Ensembl gene_id.

symbol: Ensembl gene symbol.

gene_biotype: Ensembl biotype annotation.

Direction: Concordant or Discordant fold-change between skin and PBMCs.

TISSUE FoldChange: Fold-change detected in given tissue.

TISSUE qval: False-discovery rate corrected p-value in given tissue.

Weighted gene co-expression analysis: ST8, ST11, ST13, ST16, ST19, ST21

gene_id: Ensembl gene ID.

symbol: Ensembl gene symbol.

gene_biotype: Ensembl biotype annotation.

moduleColor: Color assigned to expression module in WGCNA.

Correlation: Calculated correlation between gene and trait.

pvalue: P-value for the correlation.

Shared WGCNA correlations: ST24

gene_id: Ensembl gene ID.

symbol: Ensembl gene symbol.

gene_biotype: Ensembl biotype annotation.

Trait: clinical trait the gene is correlated with.

Directionality: Concordant or discordant, depending on if the sign of the correlation was the same in each tissue.

TISSUE Correlation: the correlation value with the trait in the given tissue.

TISSUE pvalue: correlation p-value in the given tissue.

Gephi network statistics: ST25-26

gene_id: Ensembl gene ID.

symbol: Ensembl gene symbol.

gene_biotype: Ensembl biotype annotation.

trait: clinical trait the gene is correlated with.

traitcount: number of traits the gene is correlated with (may be more than 1).

Degree – eigencentrality: network statistics calculated in Gephi Network Overview.

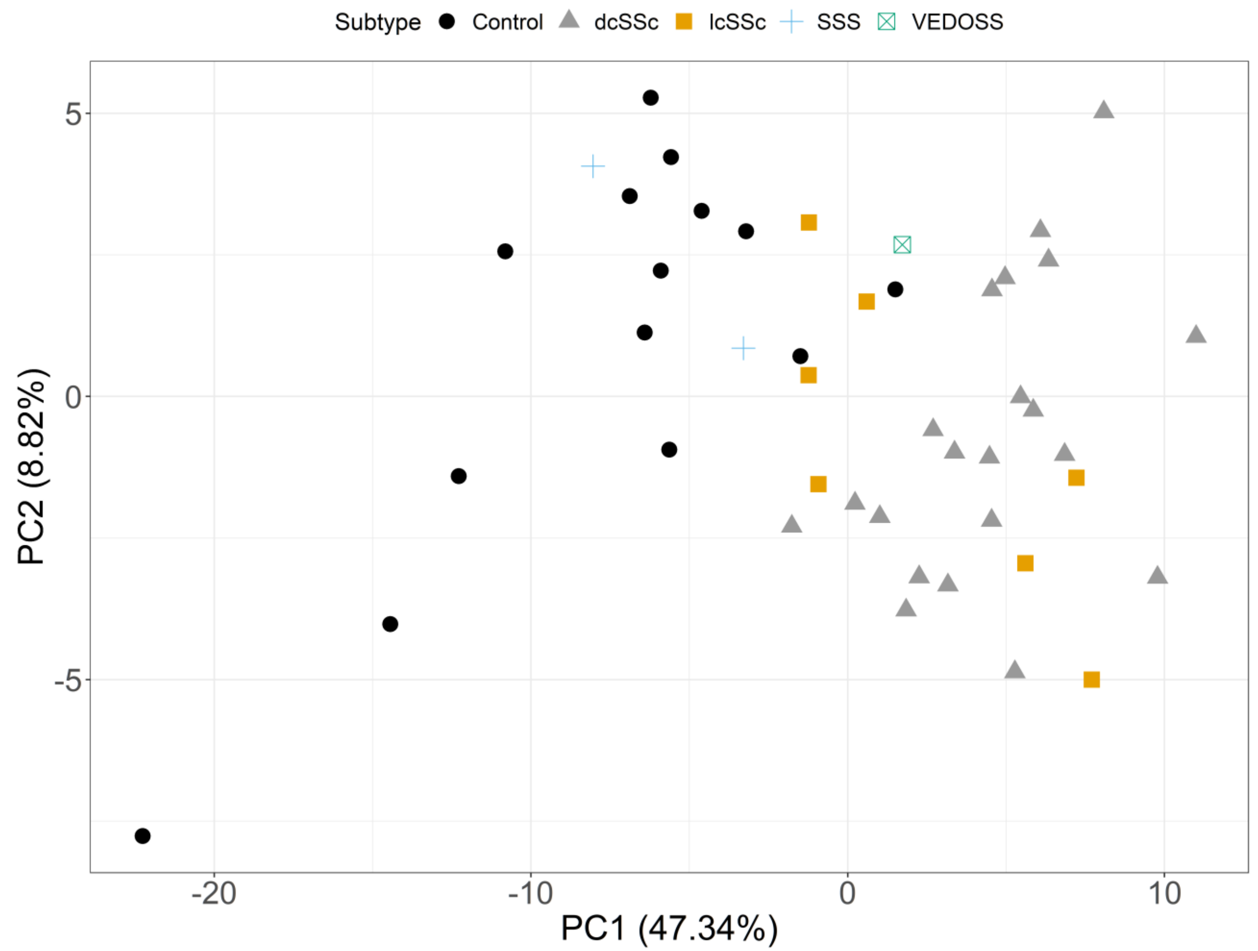


**Figure S1 – PBMC transcriptome PCA fails to distinguish SSS and VEDOSS from controls**

PCA of control and systemic sclerosis samples from all subsets, including follow-up biopsies for top 100 differentially expressed PBMC genes. The main case/control differences are in the first principal component (47%), which mostly separates the controls from SSc samples. The SSS, VEDOSS, and some limited cutaneous samples are decidedly control-like. Other limited cutaneous samples are indistinguishable from diffuse cutaneous SSc. The primary PBMC signal was in type 1 interferon signaling. In that respect, the SSS and VEDOSS samples may have less strong type 1 interferon activation.

Abbreviations: PBMC, peripheral blood mononuclear cells; PCA, principal components analysis; SSc, systemic sclerosis; SSS, SSc sine scleroderma; VEDOSS, very early diagnosis of systemic sclerosis.

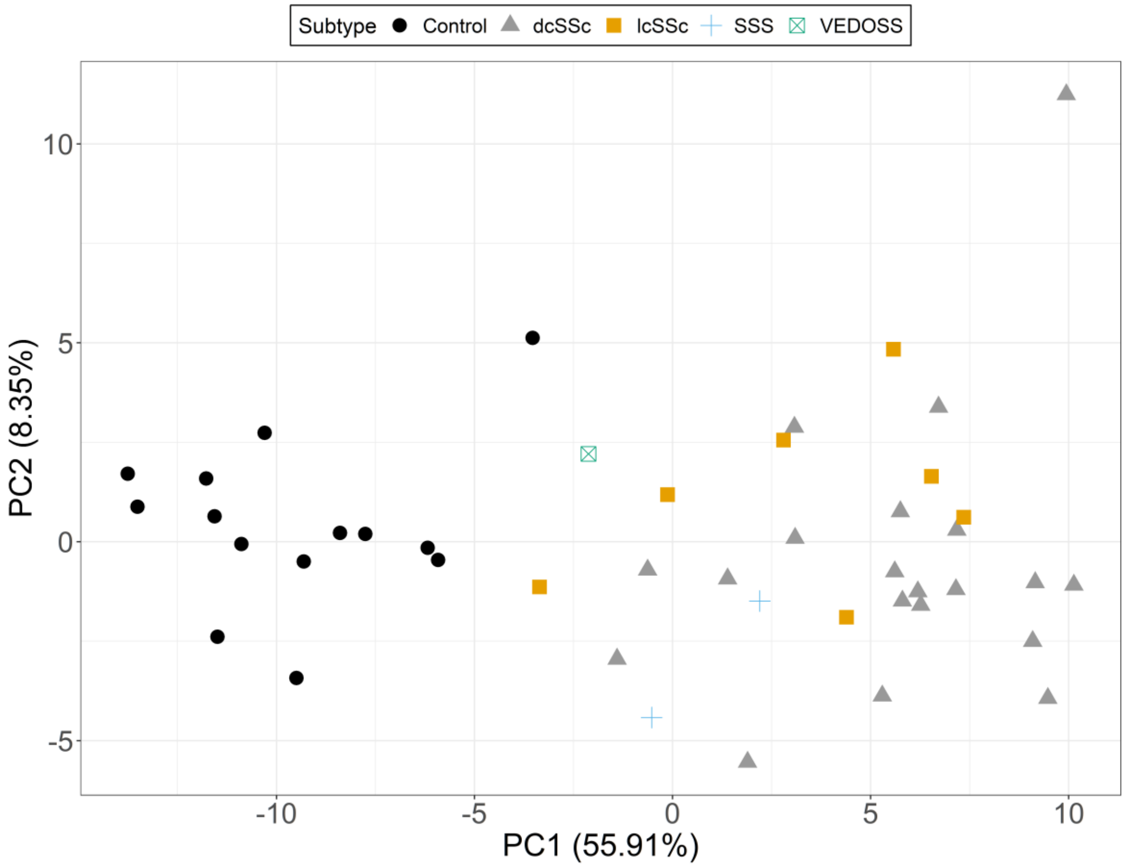


**Fig. S2 – Skin transcriptome PCA clusters SSS samples with dcSSc**

PCA of control and systemic sclerosis samples from all subsets, including follow-up biopsies for top 100 differentially expressed skin genes. The first principal component (55.9% variance explained) cleanly separates control samples from all SSc samples. In this case, the VEDOSS sample and some lcSSc samples are more control-like, but still do not cluster amongst the controls. Unexpectedly, the SSS samples cluster along with the dcSSc samples, despite not having overt dermal fibrosis. It is unclear if this is an effect of SSS patients not receiving a crucial trigger, or having some intrinsic resistance to dermal fibrosis.

Abbreviations: PCA, principal components analysis; SSc, systemic sclerosis; SSS, SSc sine scleroderma; VEDOSS, very early diagnosis of systemic sclerosis; lcSSc, limited cutaneous systemic sclerosis; dcSSc, diffuse cutaneous systemic sclerosis.
